# Are Maximum Yields sustainable? Effect of intra-annual time-scales on MSY, stability and resilience

**DOI:** 10.1101/2022.09.26.509503

**Authors:** Antoine Ricouard, Sigrid Lehuta, Stéphanie Mahévas

## Abstract

The concept of Maximum Sustainable Yield (MSY) have been lying at the core of the theory of sustainable harvesting a fishery for decades and have become a key reference point for many fishing administrations, including the European Union. However, the existence of a MSY relies on the stability of a population equilibrium. This hypothesis, though always true in the original Schaeffer model, is still challenging in more realistic and recent population models. However, recent advances shows that fish population can exhibit complex dynamics that are ill described by the classical theory. In particular, processes occurring at intra-annual time scales can affect the stability of a population equilibrium even in a strictly single species case. Associated to stability, the resilience of the equilibrium (defined as an inverse return-time following a perturbation) also matters in a management purpose. Here, we introduce an analytical single population model in discrete time with a monthly time-step allowing temporal distinction between maturation and recruitment with density-dependent mortality and fishing exploitation. We show that, thanks to an appropriate population structure, we can easily derive inter-annual population equilibrium, and study their resilience and stability properties. Then, we show that under classical hypothesis concerning density-dependence, equilibrium stability is not guaranteed and that MSY can, in theory, be associated to unstable or low resilient states. However such destabilisation seems unlikely with realistic sets of parameters. Finally, a numerical illustration for sole (*Solea solea*) of the Bay of Biscay suggests that the value of MSY was sensitive to maturation period whereas viability, stability and resilience was more sensitive to timing of recruitment. The value of *F_MSY_* appeared robust to uncertainty concerning maturation and recruitment. We conclude by saying that even if the risk of destabilisation is low for real populations, the risk of decreased resilience near the border of extinction should be cared of.

## 1. Introduction

Numerous examples of marine population collapses (Mullon et al., 2005; Pauly et al., 2005) led to the progressive recognition that world fisheries were exhaustible and that fishing could affect deeply the abundance of marine populations (Pauly et al., 2002). This concern, along with the will to maximize profits out of fisheries exploitation, fostered the development during the *XX^th^* century of a theory of sustainable harvesting of a population. The idea that there is an optimal level of fishing effort emerged after pioneer works of Russell (1931), Hjort (1933) and Graham (1935), and led to the formalisation of the concept of Maximum Sustainable Yield (*MSY*) by Schaefer (1954).

Although early criticised (Larkin, 1977), this concept was highly successful amongst several fishing administrations worldwide (Mace, 2001; Mesnil, 2012), including the European Union which set the goal that all stocks reach levels of biomass compatible with the production of MSY by 2020 (European Union, 2013). For Finley (2009), however, this institutional success is more explained by its political implications than by the scientific strength of the concept in itself.

In practice, *MSY*-based management have evolved from a target point to be attained at all costs to a target range around the maximum taking into account uncertainties and allowing room to consider other management objectives or ecosystem aspects (Hilborn, 2010; Rindorf et al., 2017). In particular, the effect of uncertainty on several model inputs on the *MSY* have been largely studied (Zheng et al., 2019) and serves as a basis for fisheries advice (ICES, 2015). However, the sustainability of the *MSY* is still not clearly established and the effects of uncertainty of inputs on sustainability of this reference point have been little explored.

The concept of sustainability is ubiquitous in the policy realm but lack clear definition which does not always match with those employed by scientists (Hilborn et al., 2015; Donohue et al., 2016). In the classical understanding of sustainability in fisheries as defined by Quinn and Collie (2005), and which correspond to the early developments of harvesting theory, “sustainable” is equivalent to “asymptotically stable” with the use of equilibrium models. Indeed, in the original model of Schaefer (1954) the harvested population is described by a single differential equation, admitting a unique and stable positive equilibrium. This is in line with the ancient conception that, neglecting random fluctuations due to external factors, populations tend to stabilise around an equilibrium value constrained by their environment. It must be stressed out that in this framework, any population level can be considered as sustainable as long as it is positive and MSY is a natural target for fishing management (Quinn and Collie, 2005). Even if the perception of sustainability have evolved and now includes a large number of metrics (Quinn and Collie, 2005; Hilborn et al., 2015; Donohue et al., 2016), the equilibrium-based concept of MSY, as a target or as a threshold (Mace, 2001), remains at the core of fishing management policies.

However, stability of exploited populations dynamics is not guaranteed. Hsieh et al. (2006) showed empirically that increased fishing pressure had a destabilising effect on populations in the sense that it tends to increase abundance fluctuations. There is a growing debate concerning the processes implicated in this destabilisation (Shelton and Mangel, 2011; Sugihara et al., 2011; Rouyer et al., 2012) but Anderson et al. (2008) argued that increased fluctuations were probably due to intrinsic dynamical effects associated to changes in life-history parameters (*e.g*. intrisic growth rate) in response to fishing.

Beside the binary opposition between stable and unstable attractors in population dynamics, the conceptually neighbouring notion of resilience, defined as an inverse return-time to the equilibrium Pimm (1984), have important management implications and is arousing a growing interest in ecological literature (Grimm and Calabrese, 2011). Key questions related to this notion are (i) whether or not ecological systems are likely to recover from a perturbation fast enough so that the equilibrium-based approach remain meaningful, and (ii) how exploitation and management are likely to affect this recovering capacity. Several theoretical studies have thus recently explored the effect of harvesting on resilience in relation with yields in single structured population models (Lundström et al., 2019), in prey-predator systems (Tromeur and Loeuille, 2017) or in tri-trophic food-chains Kar et al. (2019).

Many fish populations characterised by birth-pulse growth with well distinct cohorts (Laurec and Le Guen, 1981) are straightforwardly modelled by the use of stock-recruitment relationships (Ricker, 1954). In this kind of models, recruitment is a discrete event and is well represented by difference equations. Such models are known to allow cyclic and chaotic dynamics even for a single population (May, 1975) and lead to very complex dynamics (Tang and Chen, 2002). This stresses out the importance of the mathematical formalism employed and suggests that stability properties should not be taken for granted when deriving reference points such as MSY.

A drawback of the use of stock recruitment relationships is the fact that they synthesise in a single equation a large number of life-history processes occurring at the youngest stages of individuals’ life (Needle, 2002). In Ricker (1954) pioneer work for example, maturation and recruitment are confounded. However, life-history features such as maturation delay (Cole, 1954; Tuljapurkar, 1990; Koons et al., 2008) can have a large impact on the population dynamics. Timing and duration of density-dependent processes, including at time-scales shorter than one year, are also likely to have important consequences (Ratikainen et al., 2007). For example, timing of seasonal harvesting is known to affect the value of MSY (Kokko and Lindström, 1998; Xu et al., 2005) and the stability of population equilibrium (Cid et al., 2014). However, intra-annual processes are generally ignored in practice when deriving reference points for harvested fish populations. This constitutes in itself a specific form of uncertainty in models, which is likely to have important management implications (Ratikainen et al., 2007).

In this study, we consider that stability *sensu stricto* and resilience are key properties of sustainability. Here, we propose a theoretical model of a single harvested population submitted to birth-pulse growth and interstage density-dependence of juveniles. The latter proceeds by cannibalism (or other induced mortality) of immature individual by mature ones, which is well documented in a number of fish populations (Smith and Reay, 1991) including some of importance for exploitation such as cod (Bogstad et al., 1994; Uzars and Plikshs, 2000), and is known to be a major source of instability in populations (Ricker, 1954). Our aim is to use this model to investigate the consequences of the description at intra-annual time-scale of two critical processes, namely (i) maturation and (ii) recruitment, on long-term yields and their sustainability. A particular attention is given to the effect of these processes on the MSY. The first process investigated, maturation, is a purely biological process of critical importance that could not be controlled by management. Maturation is tightly linked to density-dependence, which is generally supposed to affect strongly immature individuals (Ricker, 1954; Rindorf et al., 2022), and to reproduction. It is subject to uncertainty, given that knowledge for real species is generally available on a yearly basis only (ICES, 2018). On the contrary, the second process, recruitment, defined as the young fish arrival in the exploited portion of the population, can be considered as a management control variable in the sense that is is partially dependent on fishing equipment and behavior (Laurec and Le Guen, 1981). As being related to exploitation, it is likely to have consequences on long-term yields and MSY.

This model and the whole set of hypotheses are presented in section 2. We then make the link between the monthly and yearly dynamics in section 3, by showing that the dynamics can be represented by a first order difference equation system. It is then possible to compute an inter-annual equilibrium and to study his property within the classical dynamical system theory framework. The modelling approach exposed in these sections is one the main points of the present paper. In particular, we stress on the idea that if we develop here an application constrained by a number of assumptions that limit its generality, we opened the door to the development of alternative models with varied assumptions but based on the same approach. In section 4, we give some details concerning the equilibrium properties of interest. We then apply our model to a real population with data for the Bay of Biscay sole (*Solea solea*). These applications are presented in section 5. Finally, our findings are discussed in section 6.

## 2. Models & analysis

### 2.1. Construction of a properly-structured population model with a monthly time-step

#### 2.1.1. Population & time structure

Throughout this study, we model the monthly discrete dynamics of a marine population exploited by fishing from a particular biological development stage (recruited individuals). The population is described with a structured abundance at each month *t*, represented by the vector <u>*N*</u>(*t*) with *t* = 1, …, +∞. Let us assume that in their life-time, individuals go through three structuring events: (i) reproduction (*i.e*. production of immature individuals from mature ones), (ii) maturation (*i.e*. transformation of immature individuals into mature ones) and (iii) recruitment. The latter is defined (Laurec and Le Guen, 1981) as the entry of individuals into the exploited portion of the population. For sake of simplicity we assume that only mature individuals are exploited, *i.e*. maturation occurs before recruitment.

Contrarily to annual models where the life-cycle events undergone by individuals are assumed simultaneous, we assume here that new-born individuals mature and recruit after a fixed number of months noted Δ*_mat_* and Δ*_rec_* respectively (as represented on time arrow in figure 1). Between two successive time-steps *t* and *t* + 1, events happening in individuals lives depend on their exact age *δ* (in months) in comparison with Δ*_mat_* and Δ*_rec_*. Immature individuals (*δ* ≤ Δ*_mat_*) are unable to reproduce, are not exploited and are assumed to undergo each month *t* a density-dependent mortality from mature individuals. Mature individuals (*δ* > Δ*_mat_*) are able to reproduce and are assumed to undergo each month a constant natural mortality. Recruited individuals (*δ* > Δ*_rec_*) undergo an additional fishing mortality and contribute to yields.

**Figure 1:**
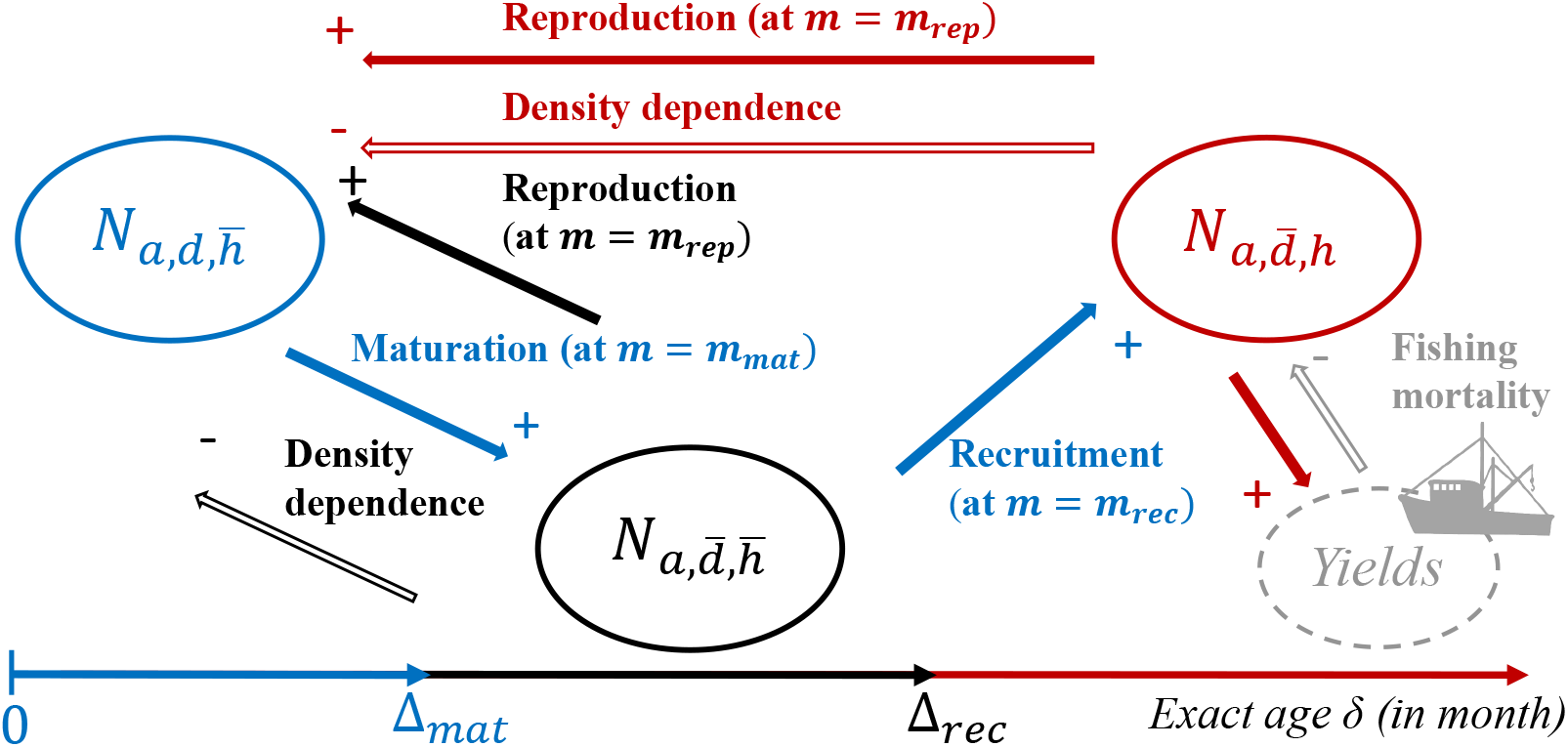
Conceptual view of the interactions between individuals and their influence on yields. Index {*a* = 0, …, > *a_rec_*} stands for round age in years, index 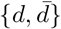 stands for density-dependence and index 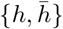 stands for accessibility to fishing. Three groups must be considered depending of exact age *δ* (in months) of individuals : *δ* < Δ*_mat_* (in blue), Δ*_mat_* ≤ *δ* < Δ*_rec_* (in black) and Δ*_rec_* ≤ *δ* (in red). Depending on Δ*_mat_* and Δ*_rec_*, the relative duration of each fraction of the population will vary and have consequences on yields and their sustainability.

As represented in figure 1, the result of this distinction between immature, mature, unharvested and harvested individuals is that three homogeneous groups of individuals (represented in different colors on the figure) interact with each other in different ways. Depending on the value of Δ*_mat_* and Δ*_rec_*, the relative importance of each process is expected to differ. First, density-dependence is expected to have a larger effect on the overall dynamics if Δ*_mat_* is large. The relative position of Δ*_rec_* to Δ*_mat_* controls the amount of time spent by mature individuals in the unspoiled situation characterized by a higher survival rate than immature and recruited individuals. The longer this duration time (Δ*_rec_* – Δ*_mat_*) is, the more numerous mature protected individuals are, hence fostering reproduction. However it also affects the number of immature individuals through the capacity of fishing to modulate the effect of natural compensation within the population by removing cannibalistic adults. Indeed, individuals of age Δ*_mat_* ≤ *δ* < Δ*_rec_* do not undergo fishing mortality but exerts density-dependent mortality on immature ones.

In order to investigate the effect of the intra-annual timing of population structuring events ((i),(ii) and (iii) detailed above) on the long-term dynamics, we define the annual cycle as the repetition of a reproduction event. For simplicity reasons and without losing in genericity, we assumed that reproduction occurs at month *m_rep_* = 12. Δ*_mat_* and Δ*_rec_* can be expressed as:

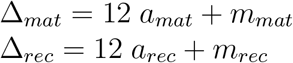

where *m_mat_* and *m_rec_* are the respective months where maturation and recruitment happen each year, and *a_mat_* and *a_rec_* are the respective number of whole years before these events happen in individuals’ life. We have 1 ≤ *m_mat_* ≤ *m_rep_* = 12 and 1 ≤ *m_rec_* ≤ *m_rep_* = 12. Assuming Δ*_mat_* ≤ Δ*_rec_*, we also have *a_mat_* ≤ *a_rec_*.

To switch from the time step *t* to a calendar time, We also define a bijective function *f* which to each time-step *t* associates a value of month m and year *y*:

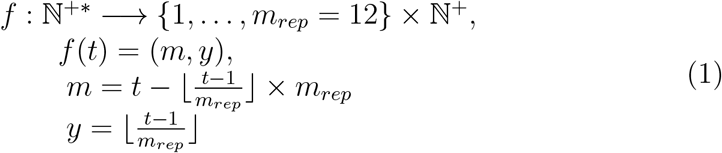

where 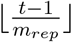 stands for the integer part of 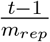, and *m_rep_* is the month at the end of which reproduction happens.

We will then denote *<u>N</u>*(*m, y*) the vector of abundance at the end of month *m* of the year *y*.

One of the main originalities of our model lies in the choice of the structure of the population *<u>N</u>*(*t*) according to successive properly defined stages. Thanks to this particular structure, we will be able to perform easily the equilibrium and stability analysis in the next step. The stages are defined by the round age *a* in year (expressed as an integer number of whole years lived by individuals, as opposed to exact age *δ* in months), the maturity (characterised by constant instead of density-dependent natural mortality) and the accessibility to fishing. Individuals are arranged into *a_rec_* + 2 modalities associated to round age *a*: {0, 1, …, *a_rec_*, > *a_rec_*} (where > *a_rec_* stands for individuals of age strictly greater than *a_rec_*), two modalities associated to maturity (density-dependant vs. independent): 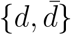 and two modalities associated to fishing (harvested vs. unharvested) 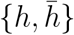.

It follows from our set of assumptions that, for all *t*:

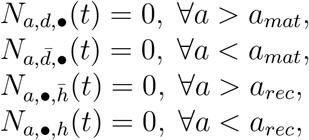

where *N_a,d•_*(*t*), 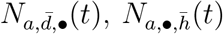, *N_a,•,h_*(*t*) stands for the abundance of respectively immature, mature, harvested and unharvested individuals of round age *a* at time *t*.

Hence, we can write the vector of the structured population:

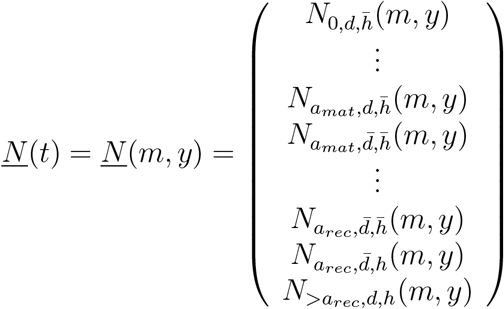

Figure 2 gives a schematic view of the complete population and time structure of our model.

**Figure 2:**
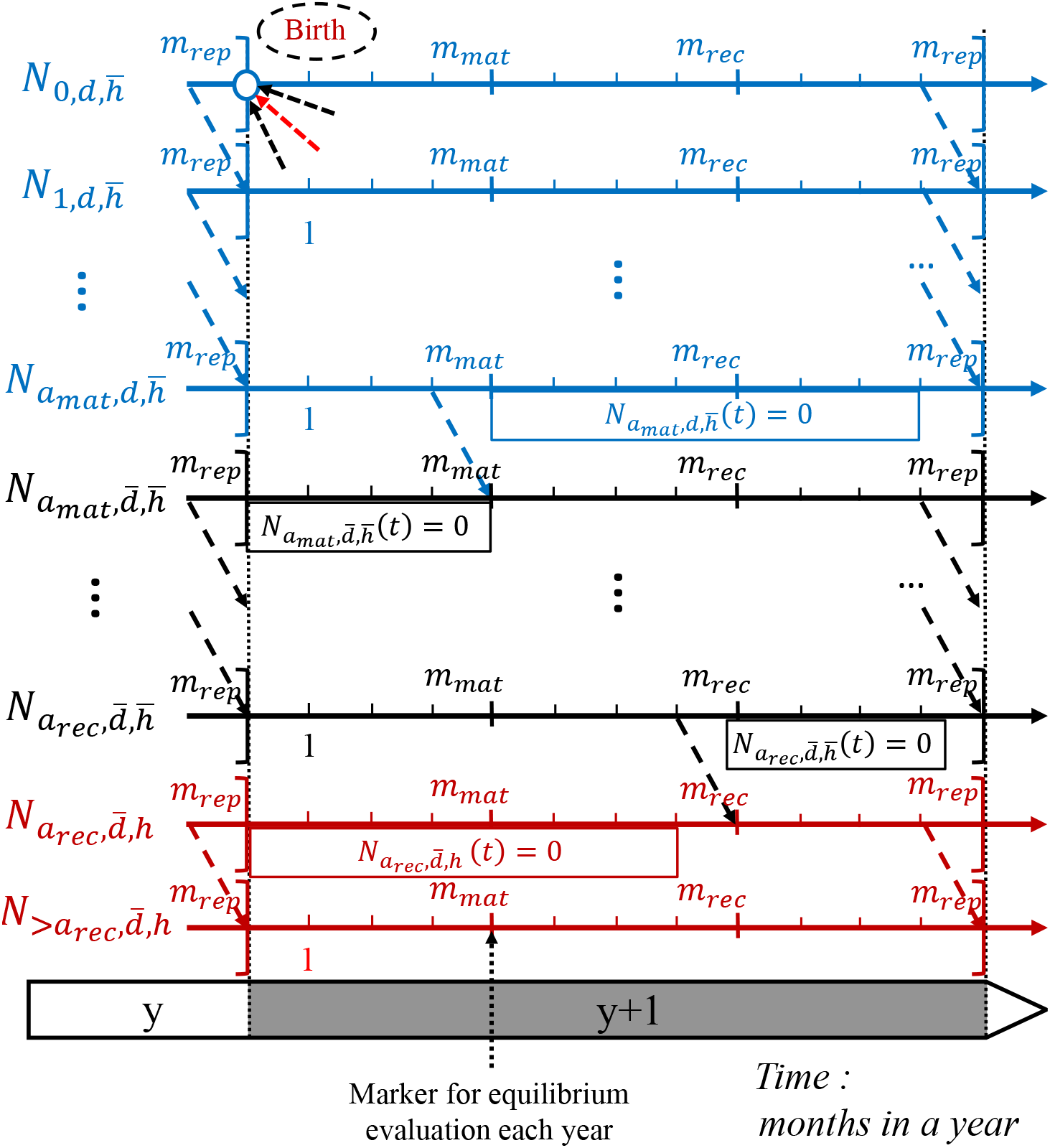
Conceptual view of the structured population and its dynamics represented in our model. Graduations are in months, and year succession is represented by the large arrow at the bottom, for all year *y*. The blue circle with dashed red and black arrows stands for the production of individuals of age 0 (“Birth”) at month *m* = *m_rep_*. Dashed coloured arrows stand for class changes (*i.e*. aging, maturation or recruitment). Empty rectangles below axes mean that some classes are always empty during part of the year because of a class change happening during the year. Each month, individuals undergo a defined mortality as shown in figure 1. The position of month *m_mat_* is indicated as being the month at which inter-annual equilibrium research is performed, for all year *y*.

#### 2.1.2. Description of the monthly dynamics

As outlined above, the two modalities associated to maturity 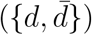 and to fishing accessibility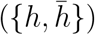 have implications in terms of mortality.

We model explicitly a density-dependent mortality of immature individuals at each time step, assuming after Ricker (1954) that mortality of immature individuals increases with the number of mature individuals. For the sake of simplicity, let the mortality of immature individuals between *t* and *t* + 1 be:

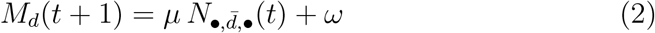

where *μ* and *ω* are two positive constants and 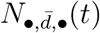 is the sum of mature individuals of all ages accessible to fishing at time *t*. The derived survival rate of immature individuals is expressed as: *S_d_*(*t* + 1) = *e*^−*M_d_*(*t*+1)^.

Let the natural and fishing mortalities of mature individuals between time-steps *t* and *t* + 1 be two positive constants 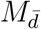 and *F*. The derived survival rates for mature individuals, depending on if they are harvested or not, are expressed as 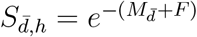 and 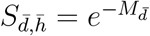.

For each *t*, switching from *t* to (*m, y*) the abundance of the population can be analytically described from the abundance at time *t* – 1. For class 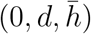, we have for all *y*:

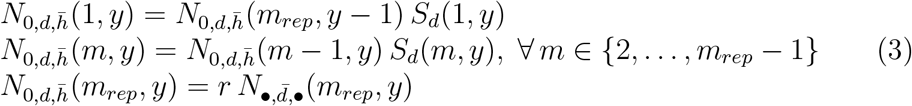

where *r* is mature individuals’ fecundity and 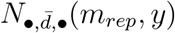 is the sum of mature individuals at the end of month *m_rep_* of year *y*.

For classes 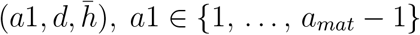, we have for all *y*:

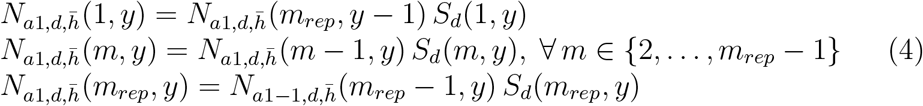

For classes 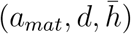 and 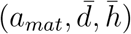, we have for all *y*:

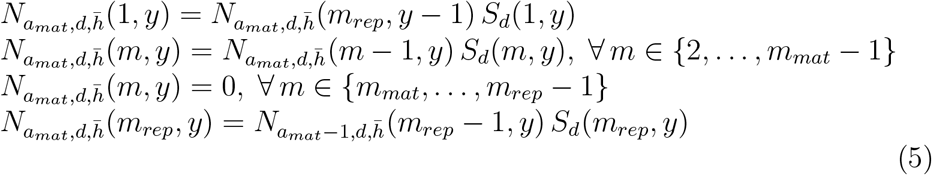

and

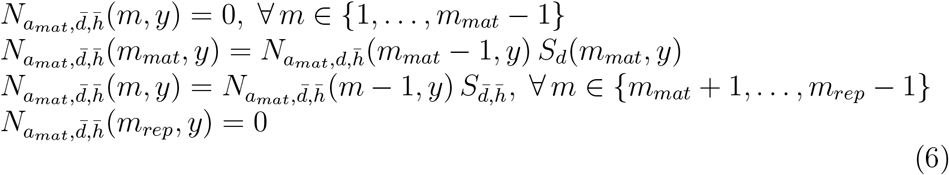

Notice that, for all *m*, only one of these last two classes takes non-zero values.

For classes 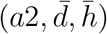, *a*2 ∈ {*a_mat_* + 1, …, *a_rec_* – 1}, we have for all *y*:

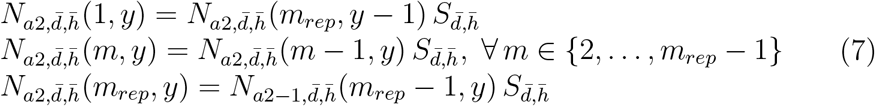

For classes 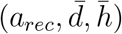 and 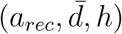, we have for all *y*:

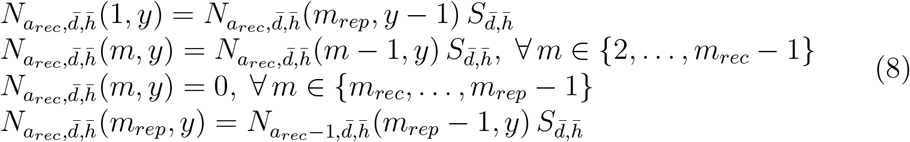

and

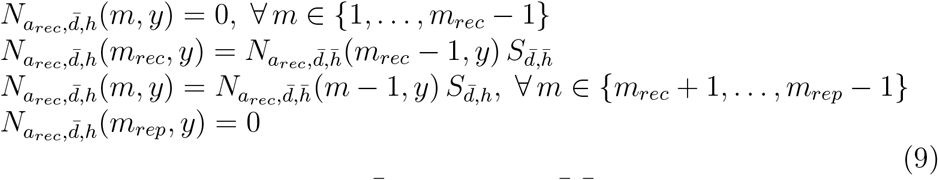

As well as for classes 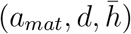 and 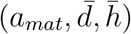, only one these last two classes takes non-zero values, for all *m*.

Finally, for class 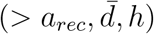, we have for all *y*:

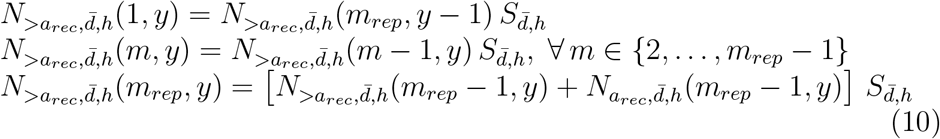

Notice that above equations were detailed for the most complex version of the model *i.e*. when 1 < *a_mat_* and 1 < *a_rec_* – *a_mat_*. However, they can be easily reduced to any version of the model with 0 ≤ *a_mat_* ≤ *a_rec_* as long as Δ*_mat_* ≤ Δ*_rec_*.

This analytical model is implemented in R and reproduces correctly the discrete monthly dynamics of exploited marine population (see figure 3 for an illustration).

**Figure 3:**
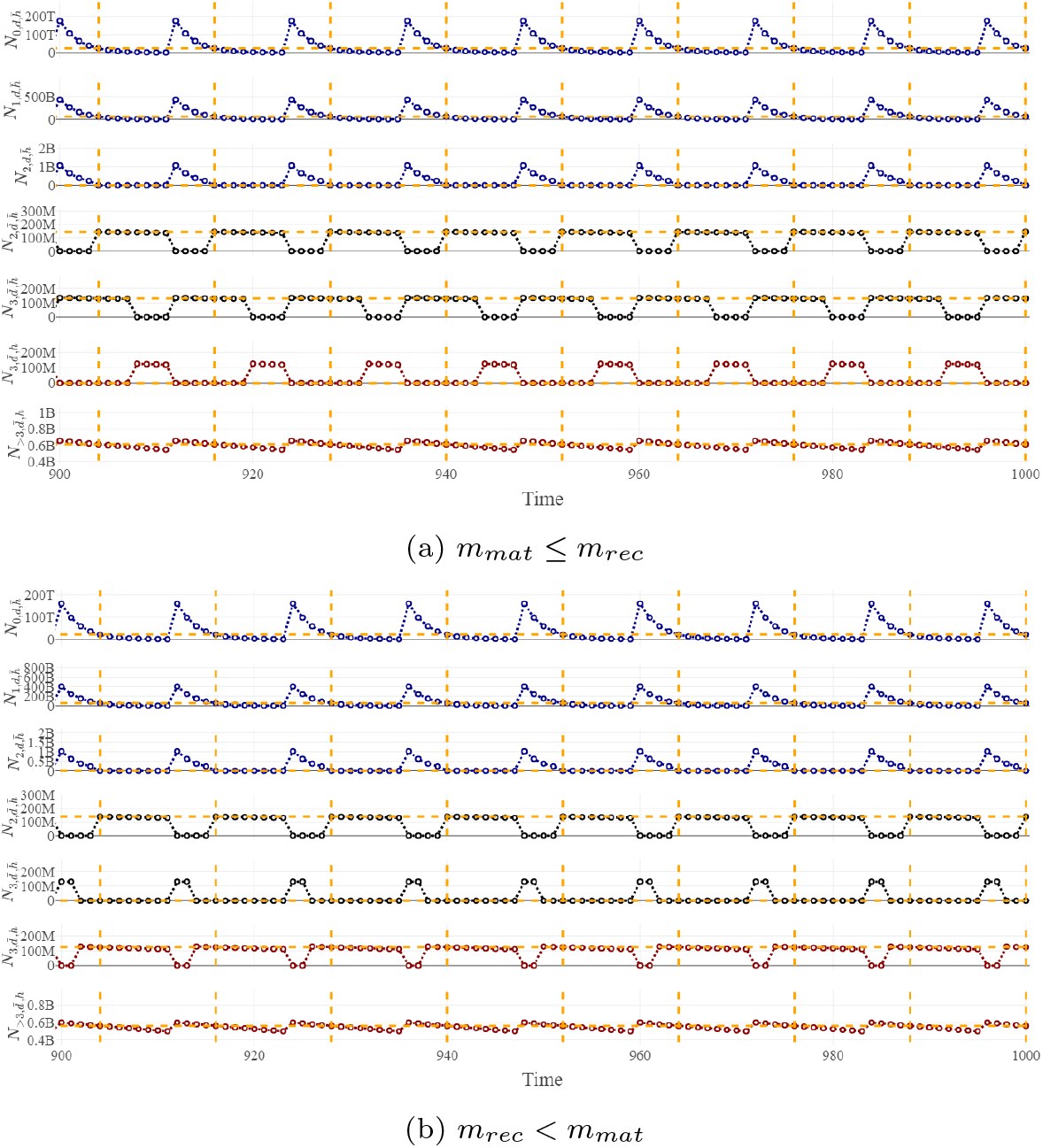
Simulated asymptotic behavior of the structured model with a monthly timestep, in both configurations (*m_mat_* ≤ *m_rec_* and conversely). Only the asymptotic abundance is plotted with month time-steps from 900 to 1000 (*t* ∈ [900; 1000]).The theoretical inter-annual equilibrium at *m* = *m_mat_* for each elements of *<u>N</u>*(*m, y*) is represented with horizontal dotted lines on each subplot. Periodic repetition of month *m_mat_* is represented by vertical dotted lines. Parameters are set for the Bay of Biscay sole (see Appendix D for parameterization details) and lags are set differently in each column: (a) Δ*_mat_* = 28 (*i.e*. *a_mat_* = 2, *m_mat_* = 4) and Δ*_rec_* = 44 (*i.e*. *arec* = 3, mrec = 8); and (b) Δ*_mat_* = 28 (*i.e*. *a_mat_* = 2, *m_mat_* = 4) and Δ*_rec_* = 38 (*i.e*. *a_rec_* = 3, *m_rec_* = 2).

### 2.2. Connecting intra-annual and inter-annual time-scales

#### 2.2.1. From a monthly dynamics to an annual dynamics

To investigate equilibrium properties of the model, we need to study the long-term evolution of the population abundance. Let *<u>N</u>*(*m, y*) be assessed at a particular arbitrary chosen month each year. Without the loss of generality, we set *m* at *m_mat_*. We will see below that the choice of *m_mat_* allows to reduce the dimension of the studied system. We therefore study the dynamics of *<u>N</u>*(*m_mat_, y*) with respect to *y*.

First, it comes from equations (5), (6), (8) and (9) that at *m* = *m_mat_*, two elements of *<u>N</u>*(*m_mat_, y*) are always empty. Depending on the the ordering of *m_mat_* and *m_rec_* we have for all *y*:

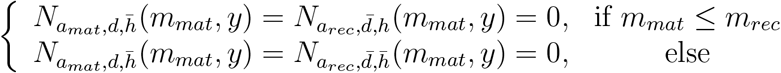

Therefore, we can always remove elements of *<u>N</u>*(*m_mat_, y*) without losing information while performing the analysis of the annual dynamics.

Then, we can derive from equations (3–10) the expression of *<u>N</u>*(*m_mat_, y*+1) for all *y*. If *m_mat_* ≤ *m_rec_*, we get the system (17), expressed in table 3. Here, we focus on this particular case but the same reasoning is feasible for the alternative case when *m_mat_* > *m_rec_* (see Appendix A for the associated developments).

**Table 1:** Time and population structure notations.

| Notation | Interpretation |
| --- | --- |
| $t$ | Model time-step (expressed in months) |
| $(m, y)$ | Time-step expressed as a calendar time (month and year) |
| $\delta$ | Exact age of individuals (in months) |
| $a$ | Round age of individuals (in years) |
| $d, \bar{d}$ | Density dependent/independent individuals (equivalent here to immature/mature individuals) |
| $h, \bar{h}$ | Harvested/unharvested individuals |
| $\Delta_{mat}$ | Lag between birth and maturation (in months) |
| $\Delta_{rec}$ | Lag between birth and recruitment (in months) |
| $a_{mat}$ | Minimum round age of matured individuals (in years) |
| $a_{rec}$ | Minimum round age of recruited individuals (in years) |
| $m_{mat}$ | Month of the year at which maturation occurs |
| $m_{rec}$ | Month of the year at which recruitment occurs |
| $m_{rep}$ | Month of the year at which reproduction occurs (by construction, we always have $m_{rep} = 12$) |
| $\underline{N}(t)$ | Vector of the structured population at time $t$ |
| $N_c(t)$ | Number of individuals of class $c$ at time $t$. The class $c$ is defined by the intersection of groups of individuals of age $a$, immature/mature individuals $(d, \bar{d})$ and harvested/unharvested individuals $(h, \bar{h})$, *e.g.* $N_{a_{mat}, d, \bar{h}}(t)$ is the number of immature, unharvested individuals of round age $a_{mat}$. |
| $\bullet$ | Union of groups, *e.g.* $(\bullet, \bar{d}, \bullet)$ stands for the class of mature individuals of all ages, whether or not they are harvested. |

**Table 2:** Model parameters and their interpretation

| Parameter | Interpretation |
| --- | --- |
| $r$ | Fecundity of mature individuals (number of eggs released in a year) |
| $\mu$ | Density-dependence factor |
| $\omega$ | Density-independent part of immature individuals' mortality |
| $M_d(t)$ | Immature individuals' mortality at time $t$ |
| $M_{\bar{d}}$ | Mature individuals' natural mortality (assumed to be constant) |
| $S_d(t+1)$ | Survival rate of immature individuals between time-steps $t$ and $t+1$ |
| $S_{\bar{d},\bar{h}}$ | Survival rate of mature, unharvested individuals between time-steps $t$ and $t+1$ (assumed to be constant) |
| $S_{\bar{d},h}$ | Survival rate of mature, harvested individuals between time-steps $t$ and $t+1$ (assumed to be constant) |
| $F$ | Fishing mortality by month (assumed to be constant) |

**Table 3:**
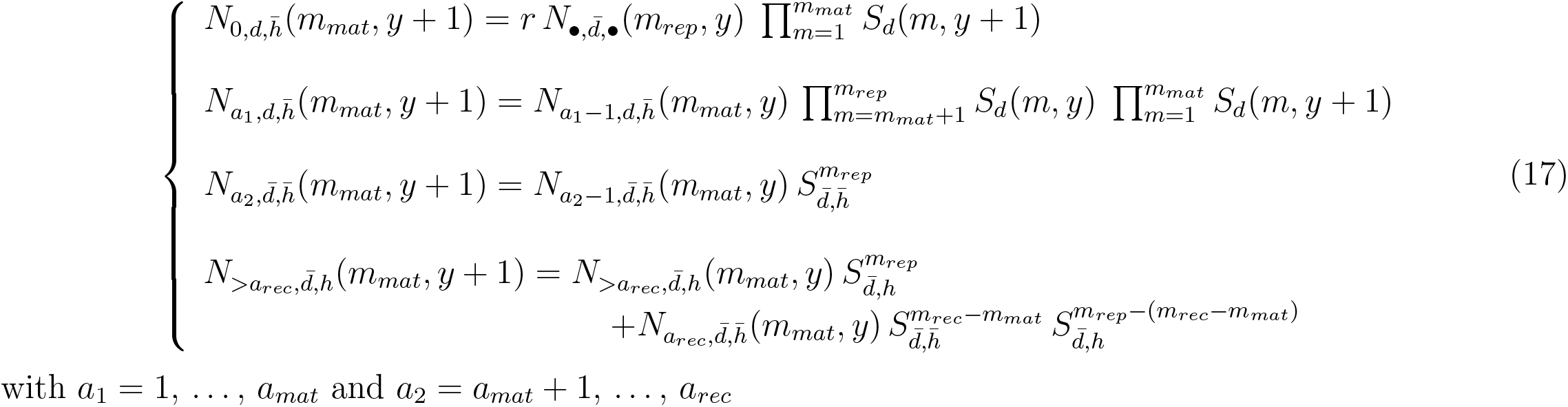
Expression of the inter-annual dynamics of *<u>N</u>*(*m_mat, y_*) when *m_mat_* ≤ *m_rec_*, for all *y*

Given some proper simplifications, 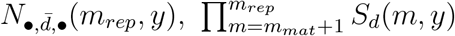 and 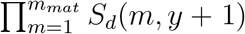 can be formulated as functions of *<u>N</u>*(*m_mat_, y*) (see Appendix B for proof) and system (17) can be expressed as a first-order difference system of dimension *a_rec_* + 2. For all *y*, we get :

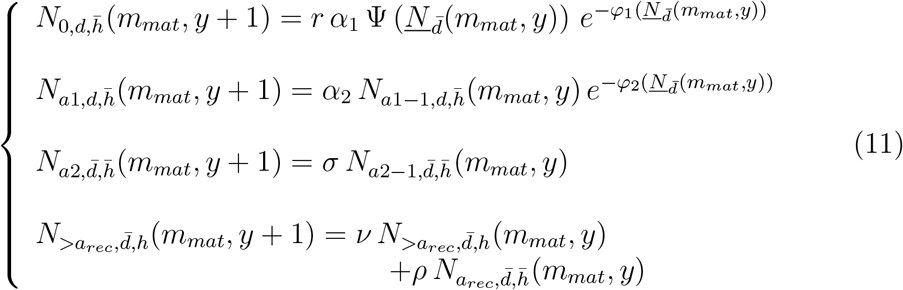

with *a*1 = 1, …, *a_mat_* and *a*2 = *a_mat_* + 1, …, *a_rec_*. 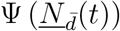 is the number of spawner and 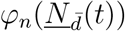 is a density-dependence function (see table 4 for detailed expressions). Moreover :

— 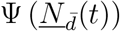 and 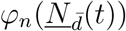 are linear combinations of 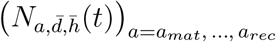 and 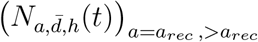.
— *α_n_, σ, v, ρ* are positive constants.

**Table 4:**
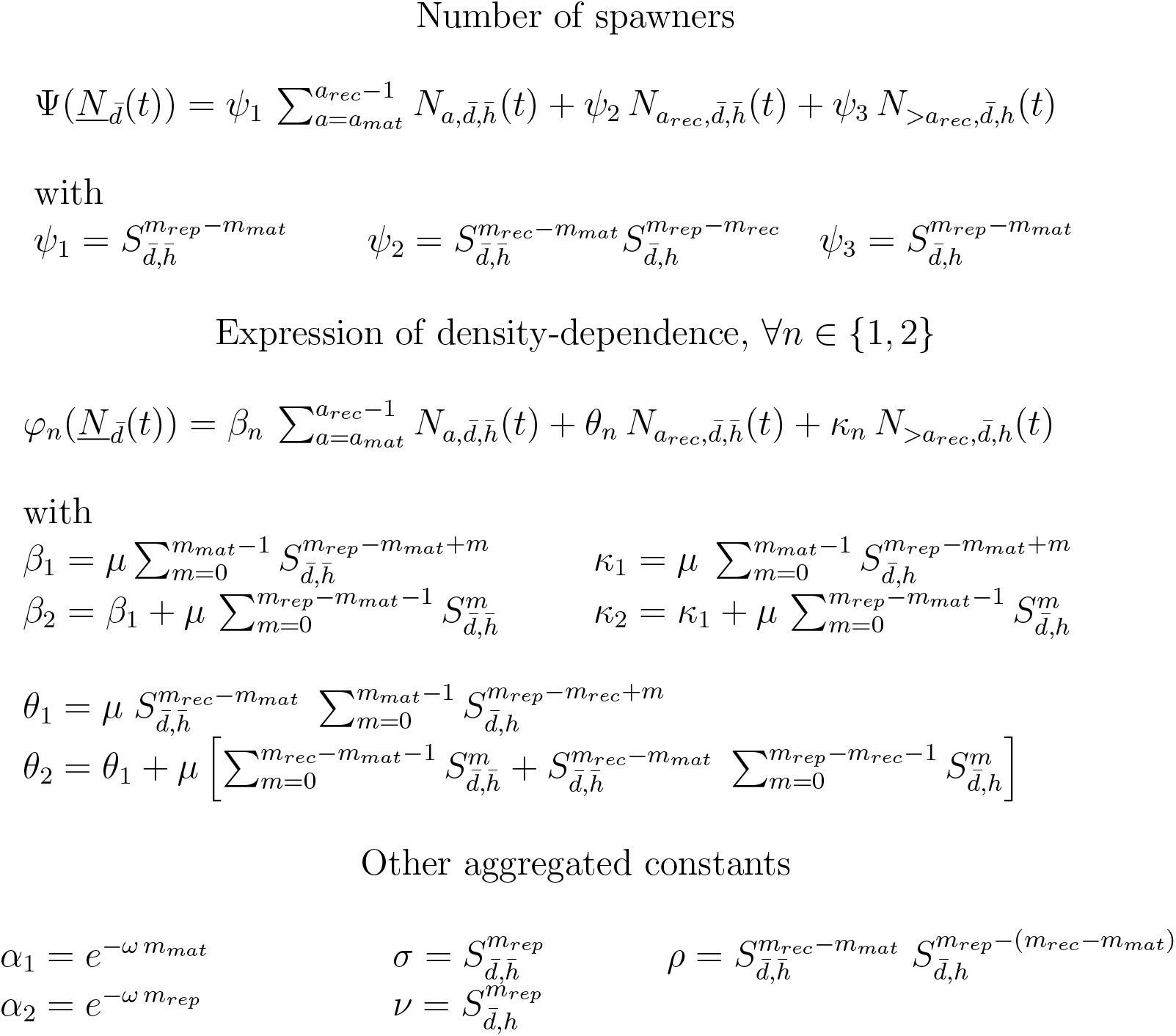
Expressions of aggregated constants and auxiliary functions used in model development. 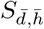 is the survival rate of mature, unharvested individuals. 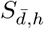 is the survival rate of mature, harvested individuals. *μ* and *ω* are the two constants in immature individuals’ mortality function (2).

At this stage, given this first order difference equations system (11) we can perform easily the equilibrium and stability analysis of the population.

#### 2.2.2. Expressions of equilibrium

In this study, we are interested in the equilibrium properties of *<u>N</u>*(*m_mat_, y*) described by system (11). *<u>N</u>*\* is an inter-annual equilibrium if and only if it verifies:

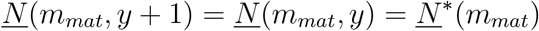

System (11) always admits only one non-trivial equilibrium which is the solution of:

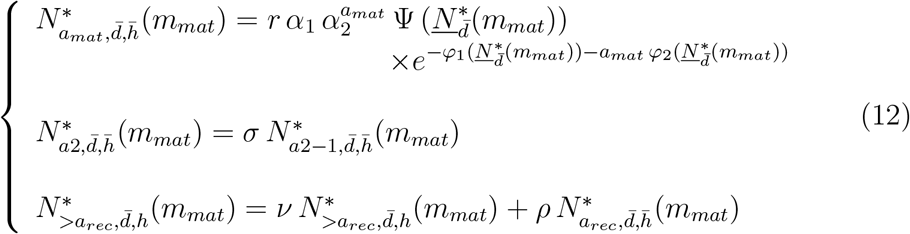

with 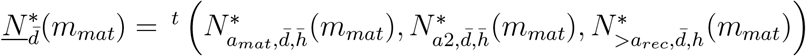 and *a*2 = *a_mat_* + 1, …, *a_rec_*;

### 2.3. Equilibrium properties : equilibrium yields, stability & resilience

#### 2.3.1. Maximum Sustainable Yield

The equilibrium yield can be straightforwardly derived from the equilibrium abundance using the classical Baranov catch equation (Baranov, 1918), given that fishing and natural mortality of mature individuals are constant :

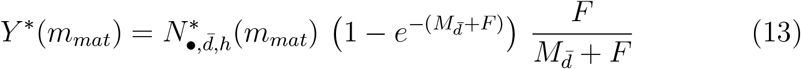

where 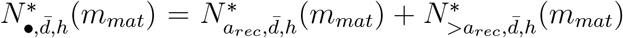 is the total number of recruited individuals at month *m* = *m_mat_* at inter-annual equilibrium.

The value of inter-annual equilibrium at any month *m* ≠ *m_mat_* can be easily computed by applying equations (3–10) to *<u>N</u>*\*. Hence we can compute the total annual yield as:

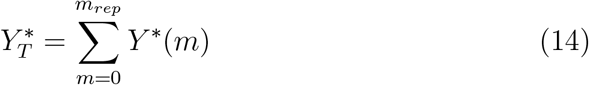

where *Y*\*(*m*) is the inter-annual equilibrium yield at month *m*.

Let us consider at present that 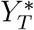 depends only on the control variable *F*. Unfortunately the expression of 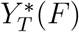 is too complex to analytically calculate the maximum of 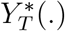 and derive the Maximum Sustainable Yield (MSY). Instead we performed a numerical optimization of 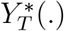 using Brent’s method (Brent, 1973) to get the value of MSY and *F_MSY_*:

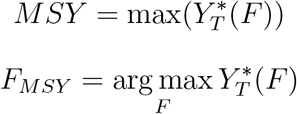

#### 2.3.2. Equilibrium stability & resilience

Computation of yields at equilibrium and their optimisation does not inform us on the stability of this equilibrium and hence on sustainability of yields. As we succeeded to express the whole dynamics of the structured population as first order difference equation system, we can perform easily the stability analysis of the equilibrium by studying the property of the Jacobian matrix of system (11). The inter-annual equilibrium *<u>N</u>*\* is locally stable if and only if:

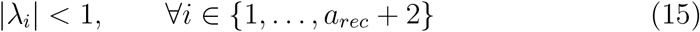

where |λ*_i_*| is the modulus of the *i^th^* eigen value of the Jacobian matrix of the system at inter-annual equilibrium. See Appendix C for the general expression of the Jacobian matrix of system (11).

Expressions of eigenvalues are expected to be too complex to be analytically tractable and interpretable especially when the system is of large dimension. We therefore calculated numerically the eigenvalues to detect stability changes using the basic “eigen” function of R software which relies on LAPACK routines (Anderson et al., 1999).

The stability properties of equilibrium depends on parameters. Hence, stability, unstability, and extinction domains (*i.e* the set of parameters for which the equilibrium is stable, unstable and lesser than zero, respectively) represent volumes in a parameter space. In particular, we pay attention of the surfaces of stability, unstability and extinction in the *r* × *F* plan. Variations along the *r* axis can represent differences of fecundity between stocks or differences in reproductive success for a same stock, whereas *F* is the main control variable when dealing with exploited systems. In such a plan, it is also possible to plot the value of *F_MSY_* if it does exist, for each value of *r*.

When no destabilisation occurs, the stability properties of the equilibrium can be more finely defined by considering the resilience of this equilibrium. *Sensu* Pimm (1984), a system is all the more resilient that the characteristic return-time to equilibrium is short. This notion is related to stability and can be studied with the same mathematical tools. Hence, in discrete time this return-time is given by (Beddington et al., 1976):

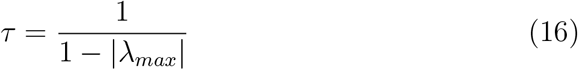

where |λ*_max_*| is the modulus of the leading eigen value of the system. The return-time is one if the system returns instantaneously to his equilibrium and infinite when the equilibrium becomes unstable.

Considering that return-time is, just as yields, a function of *F*, we can define the same way as for *F_MSY_*, a mortality *F_RMY_* for ‘Resilience Maximising Yield’ for which resilience is maximum, *i.e*. associated return-time (denoted *MiRT*) is minimum. Mathematically:

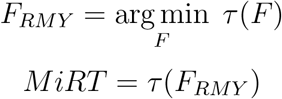

#### 2.3.3. Sensitivity of MSY, stability domain and resilience to timing of maturation and recruitment

Here, our aim is to assess the sensitivity of the equilibrium properties, namely *MSY, F_MSY_, MiRT* and *F_MiRT_* values, extinction and unstability domains (quantified by the areas under the curves *A_ext_* and *A_stab_* respectively) to intra-annual variations of maturation and recruitment, represented by parameters Δ*_mat_* and Δ*_rec_*. The first one must be considered as a biological parameter subject to epistemic uncertainty whereas the second can be considered as a control variable insofar it is, at least theoretically, possible not to catch individuals under a defined age. Values for Δ*_mat_* and Δ*_rec_* are allowed to vary in a time-span shorter than one year. Such intra-annual dynamics are generally not taken into account when computing reference points for fisheries management. Indeed, for most fish populations, information concerning maturation and recruitment are available on yearly basis only (ICES, 2018). We perform a variance-based sensitivity analysis based on 1) an experimental design and 2) sensitivity indices associated to Δ*_mat_* and Δ*_rec_* (the inputs of the model) on each metric *MSY, F_MSY_, A_ext_* and *A_stab_* (model outputs) derived from an ANOVA. Then we can compute the sensitivity index of each parameter, both for principal effect and interactions (see *e.g*. Faivre et al. (2013) for full details on the method). The experimental design is build combining all possibles values of Δ*_mat_* and Δ*_rec_* within defined ranges. These two parameters take values expanding on a time-span less or equal to one year, corresponding to realistic values for the Bay of Biscay sole (see table 5).

**Table 5:**
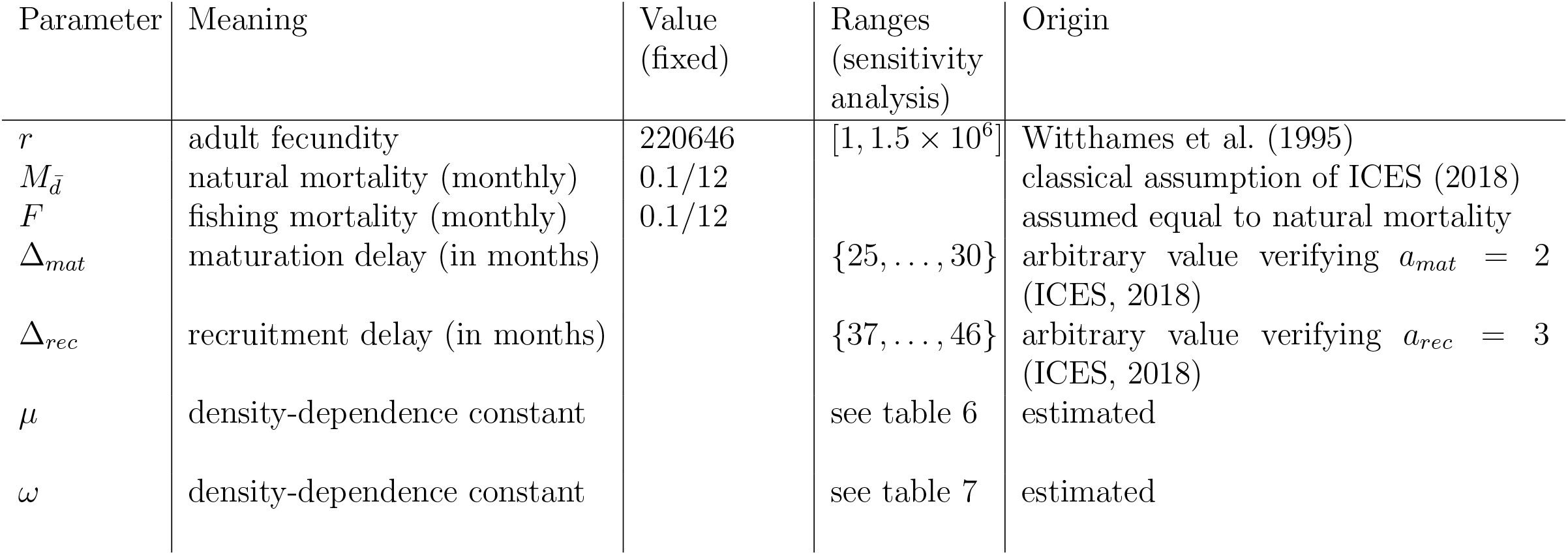
Numerical values and origin of parameters used in simulation for the Bay of Biscay sole.

| Parameter | Meaning | Value<br>(fixed) | Ranges<br>(sensitivity<br>analysis) | Origin |
| --- | --- | --- | --- | --- |
| $r$ | adult fecundity | 220646 | $[1, 1.5 \times 10^6]$ | Witthames et al. (1995) |
| $M_{\bar{d}}$ | natural mortality (monthly) | 0.1/12 | | classical assumption of ICES (2018) |
| $F$ | fishing mortality (monthly) | 0.1/12 | | assumed equal to natural mortality |
| $\Delta_{mat}$ | maturation delay (in months) | | $\{25, \dots, 30\}$ | arbitrary value verifying $a_{mat} = 2$<br>(ICES, 2018) |
| $\Delta_{rec}$ | recruitment delay (in months) | | $\{37, \dots, 46\}$ | arbitrary value verifying $a_{rec} = 3$<br>(ICES, 2018) |
| $\mu$ | density-dependence constant | | see table 6 | estimated |
| $\omega$ | density-dependence constant | | see table 7 | estimated |

## 3. Application to the Bay of Biscay sole

### 3.1. Origin of data

Our model is general and flexible enough to be applied to any exploited population as long as maturation occurs before recruitment. However, for illustration purposes and also because the complexity of our model preclude the derivation of analytical results concerning stability or yield optimisation, we performed a numerical application.

In order to get numerical simulations somewhat realistic, we parameterized our model on the ground of (i) published life-history parameters, and (ii) stock assessment data, for the sole (*Solea solea*) in the Bay of Biscay. Sole is a highly valued demersal species targeted by a number of fishing fleets in the Bay of Biscay (Vigier et al., 2022) and is subject to a stock assessment by the International Council for the Exploration of the Sea (ICES) on a yearly basis (ICES, 2018).

All but two parameters where extracted directly from literature. Numerical values of all the parameters used in simulation, with their origin and meaning are presented in table 5.

The last two parameters *μ* and *ω*, which govern density-dependence, were estimated based on ICES stock assessment results (ICES, 2018; table 7.10 p.277). The basic idea here is to fit a custom stock-recruitment relationship on data (see Appendix D for more details). To distinguish the effect of intra-annual dynamics from the effect of density-dependence, *μ* and *ω* are reestimated at each change of Δ*_mat_* and Δ*_rec_* when assessing the sensitivity of the system to these parameters. *μ* and *ω* values obtained for each pair (Δ*_mat_*, Δ*_rec_*) considered are given in tables 6 and 7.

**Table 6:** Estimation of parameter *μ* for the different combinations of Δ*_mat_* and Δ*_rec_*.

| | $\Delta_{rec}$<br>37 | 38 | 39 | 40 | 41 | 42 | 43 |
| --- | --- | --- | --- | --- | --- | --- | --- |
| $\Delta_{mat}$ 25 | $1.730 \times 10^{-11}$ | $1.644 \times 10^{-11}$ | $1.656 \times 10^{-11}$ | $1.666 \times 10^{-11}$ | $1.677 \times 10^{-11}$ | $1.686 \times 10^{-11}$ | $1.695 \times 10^{-11}$ |
| 26 | $1.909 \times 10^{-11}$ | $1.968 \times 10^{-11}$ | $1.920 \times 10^{-11}$ | $1.931 \times 10^{-11}$ | $1.94 \times 10^{-11}$ | $1.951 \times 10^{-11}$ | $1.961 \times 10^{-11}$ |
| 27 | $2.152 \times 10^{-11}$ | $2.145 \times 10^{-11}$ | $2.175 \times 10^{-11}$ | $2.160 \times 10^{-11}$ | $2.170 \times 10^{-11}$ | $2.181 \times 10^{-11}$ | $2.191 \times 10^{-11}$ |
| 28 | $2.367 \times 10^{-11}$ | $2.361 \times 10^{-11}$ | $2.359 \times 10^{-11}$ | $2.356 \times 10^{-11}$ | $2.368 \times 10^{-11}$ | $2.378 \times 10^{-11}$ | $2.388 \times 10^{-11}$ |
| 29 | $2.556 \times 10^{-11}$ | $2.547 \times 10^{-11}$ | $2.542 \times 10^{-11}$ | $2.539 \times 10^{-11}$ | $2.514 \times 10^{-11}$ | $2.549 \times 10^{-11}$ | $2.558 \times 10^{-11}$ |
| 30 | $2.722 \times 10^{-11}$ | $2.711 \times 10^{-11}$ | $2.702 \times 10^{-11}$ | $2.697 \times 10^{-11}$ | $2.694 \times 10^{-11}$ | $2.649 \times 10^{-11}$ | $2.703 \times 10^{-11}$ |
| | $\Delta_{rec}$<br>44 | 45 | 46 | | | | |
| $\Delta_{mat}$ 25 | $1.704 \times 10^{-11}$ | $1.712 \times 10^{-11}$ | $1.720 \times 10^{-11}$ | | | | |
| 26 | $1.970 \times 10^{-11}$ | $1.980 \times 10^{-11}$ | $1.989 \times 10^{-11}$ | | | | |
| 27 | $2.201 \times 10^{-11}$ | $2.210 \times 10^{-11}$ | $2.220 \times 10^{-11}$ | | | | |
| 28 | $2.398 \times 10^{-11}$ | $2.408 \times 10^{-11}$ | $2.418 \times 10^{-11}$ | | | | |
| 29 | $2.568 \times 10^{-11}$ | $2.577 \times 10^{-11}$ | $2.587 \times 10^{-11}$ | | | | |
| 30 | $2.712 \times 10^{-11}$ | $2.721 \times 10^{-11}$ | $2.731 \times 10^{-11}$ | | | | |

**Table 7:**
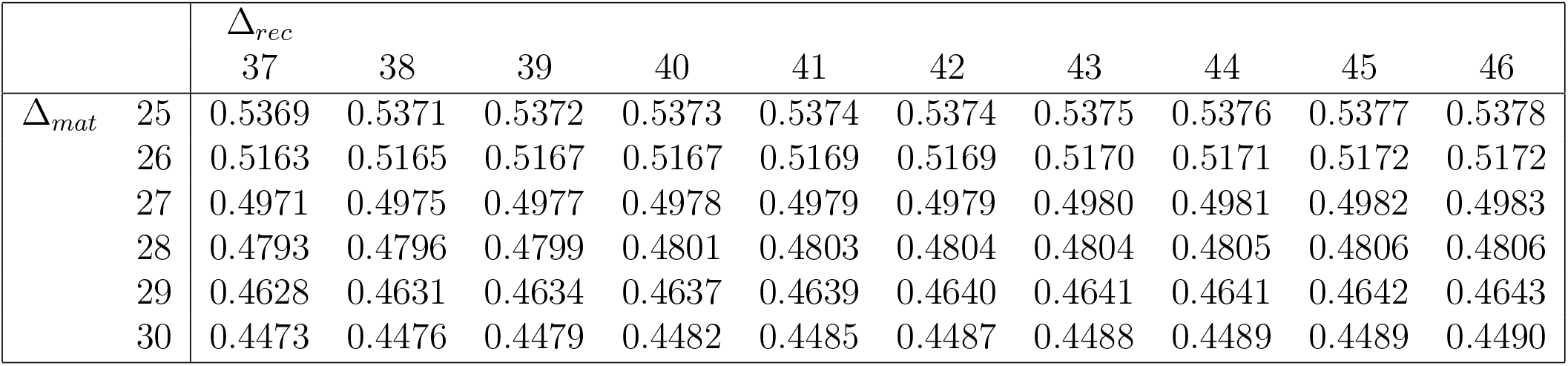
Estimation of parameter *ω* for the different combinations of Δ*_mat_* and Δ*_rec_*.

### 3.2. Monthly dynamics and inter-annual equilibrium of abundance

Once the model is fully parameterised, it is possible on the one hand to simulate the monthly dynamics with respect to equations (3–10), and on the other hand to compute the theoretical inter-annual equilibrium vector *<u>N</u>*\*(*m_mat_*). Both the abundance time series and the equilibrium abundance for each class are computed for the example of the sole of the Bay of Biscay (figure 3). In this example, the dynamics converges to a stable annual cycle represented by the inter-annual equilibrium *<u>N</u>*\*(*m_mat_*) (in dashed horizontal lines). The value of this equilibrium is expected to vary with model parameters (see Appendix E for variations of 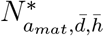 and of the sum of spawners for Bay of Biscay sole as a function of *F*, Δ*_mat_* and Δ*_rec_*).

### 3.3. Sensitivity of MSY to timing of maturation and recruitment

The equilibrium yield of the sole of the Bay of Biscay is computed for a range of monthly fishing mortality *F* ∈ [0; 0.4] and is numerically optimised as a function of *F* to get the MSY. We assessed their sensitivity to Δ*_mat_* and Δ*_rec_* varying in a range shorter than 12 months, with Δ*_mat_* = 25, 26, …, 33 (*i.e*. between the first and ninth month of the year) and Δ*_rec_* = 37, 38, …, 46.

First, it appears clearly that the value of *MSY* is much more sensitive to variations of Δ*_mat_* (*SI* = 0.98) than to Δ*_rec_* whereas the position of *F_MSY_* is sensitive to both (*SI* = 0.56 for Δ*_mat_*) as shown in figure 4a and 4b. In fact, it appears that whereas *MSY* undergoes large variations when Δ*_mat_* or even Δ*_rec_* are varied, *F_MSY_* remains remarquably constant around *F* = 0.01.

**Figure 4:**
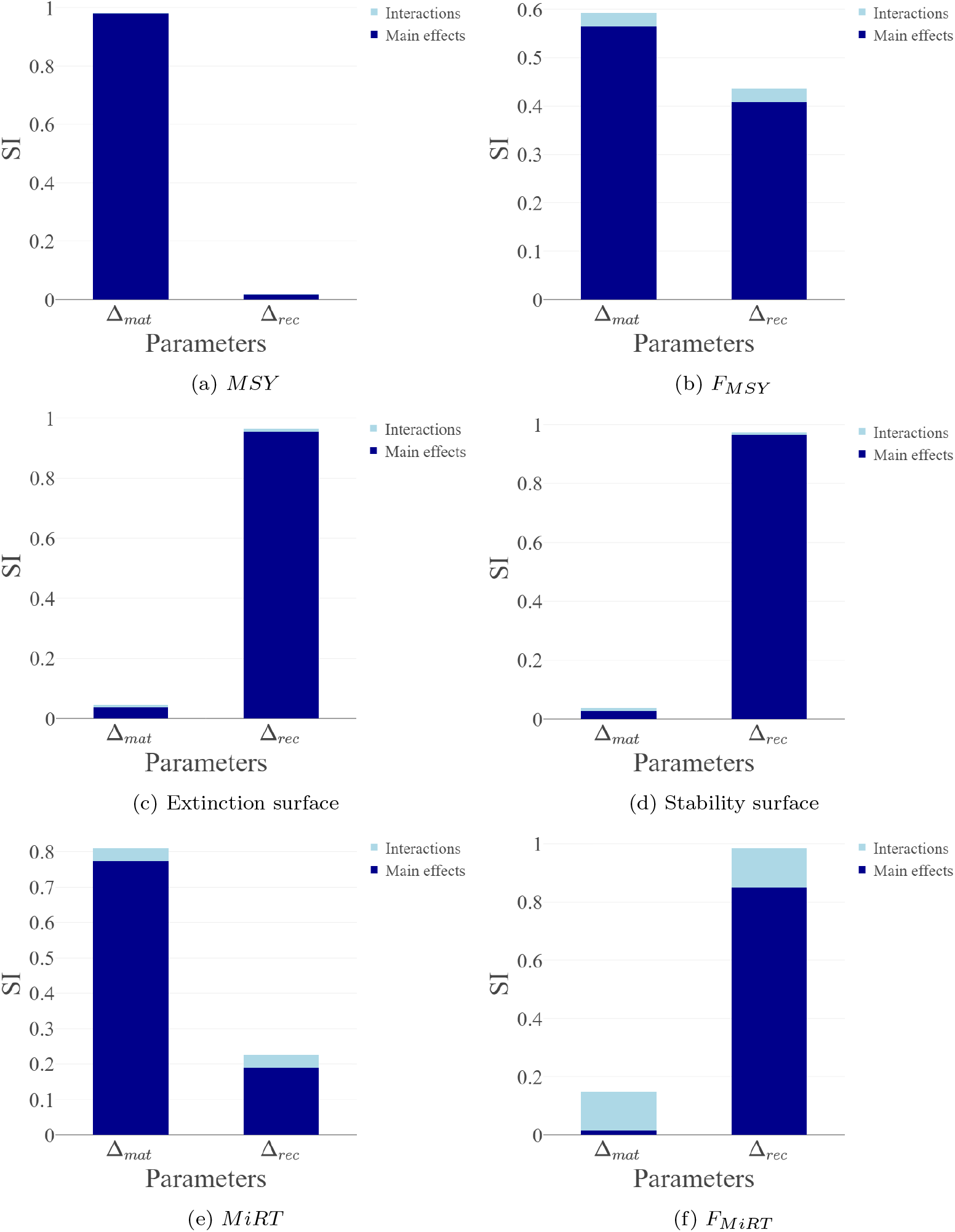
Sensitivity indices of parameters Δ*_mat_*, Δ*_rec_* for different model outputs: (a) *MSY*, (b) *F_MSY_*, (c) surface in the *r* × *F* plan as represented in figure 6 where the population goes extinct, (d) surface in this plan where the inter-annual equilibrium is unstable, (e) *MiRT*, (f) *F_MiRT_*.

When Δ*_mat_* only varies, the duration of immature phase and mature unharvested vary (respectively in blue and black on figure 1). An increase in Δ*_mat_* translates into a longer period where individuals are submitted to a large density-dependent mortality (empty arrows in figure 1) and a shorter period of protection from fishing for adults. In this case, as represented in figure 5a, MSY is maximum when Δ*_mat_* equals 25, *i.e*. when the density-dependent phase is short. An increase of one or very few months in Δ*_mat_* is sufficient to cause sharp reductions in yields and MSY. *F_MSY_*, on the contrary, increases very slightly when Δ*_mat_* increases.

**Figure 5:**
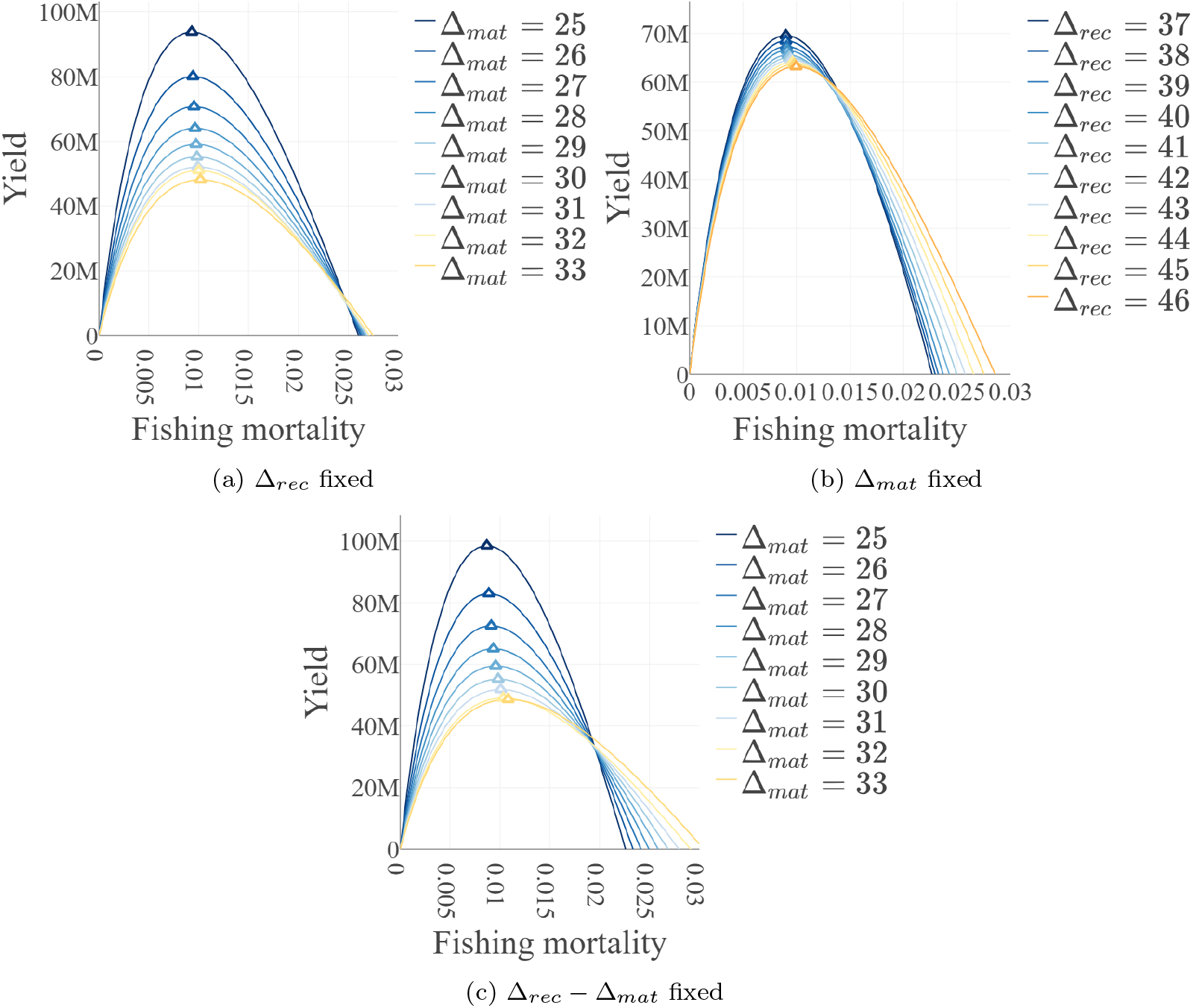
Total annual yields at inter-annual equilibrium as a function of fishing mortality (by month) and position of MSY when Δ*_mat_* and/or Δ*_rec_* vary and the system is parameterized for the sole of the Bay of Biscay. Coloured triangles correspond to MSY (numerically solved) and different colours indicate different values of Δ*_mat_* and/or Δ*_rec_* depending on the case: (a) Δ*_mat_* vary and Δ*_rec_* = 44; (b) Δ*_rec_* vary and Δ*_mat_* = 28; (c) both Δ*_mat_* and Δ*_rec_* vary and their difference is constant and equals 14 (Δ*_mat_* = 25, 26, …, 33; Δ*_rec_* = 39,40, …,47).

When Δ*_rec_* only varies and the duration of the density-dependent phase (in blue on figure 1) is kept constant. The only modification is hence on the balance between unharvested adult phase and harvested adult phase (respectively in black and red on figure 1). This translates into a modification of the ability of fishing to modulate natural regulation of the population which can be represented by the relative importance of red and black arrows on figure 1.

As shown in figure 5b, an increase in Δ*_rec_* is associated to a decrease in *MSY* but much smaller than when Δ*_mat_* is varied, as well as a slight increase in *F_MSY_* and extinction mortality (*i.e* the smallest value of *F >* 0 that brings null yields because of extinction of the population).

When Δ*_mat_* and Δ*_rec_* are varied jointly so that the mature unharvested phase (in black on figure 1) is constant in duration, the general shape of the curves obtained (figure 5c) presents properties of the first two ones. On the one hand, variations of *MSY* are large as when Δ*_mat_* only is varied, but on the other hand we get an increase in extinction mortality that was observed when Δ*_rec_* only was varied.

### 3.4. Sensitivity of stability domain to timing of recruitment and maturing

The domain of viability and of stability of the population in the *r* × *F* plan is also affected when Δ*_mat_* and/or Δ*_rec_* vary, although the second one has a much larger effect (figure 4c and 4d). Indeed, sensitivity index of Δ*_rec_* is of 0.95 for the surface of viability domain and of 0.96 for the surface of stability domain in the considered section of *r* × *F* plan, which means that most of the variance in those surfaces are explained by variations of Δ*_rec_*.

In the three cases investigated (variations of Δ*_mat_* only, Δ*_rec_* only or joint variations), population can be brought to extinction by increasing *F* or reducing *r*. Equilibrium can be destabilised by increasing *r* or increasing *F*, and *F_MSY_* always increase with *r*.

In the first case (variation of Δ*_mat_*, see figure 6a), viability frontiers are almost superposed and stability surface increases slightly when Δ*_mat_* increases. This quite surprising result suggests that a long immature phase has a stabilizing effect on the population, probably a consequence of spreading in time density-dependent processes (as illustrated in figure 1).

**Figure 6:**
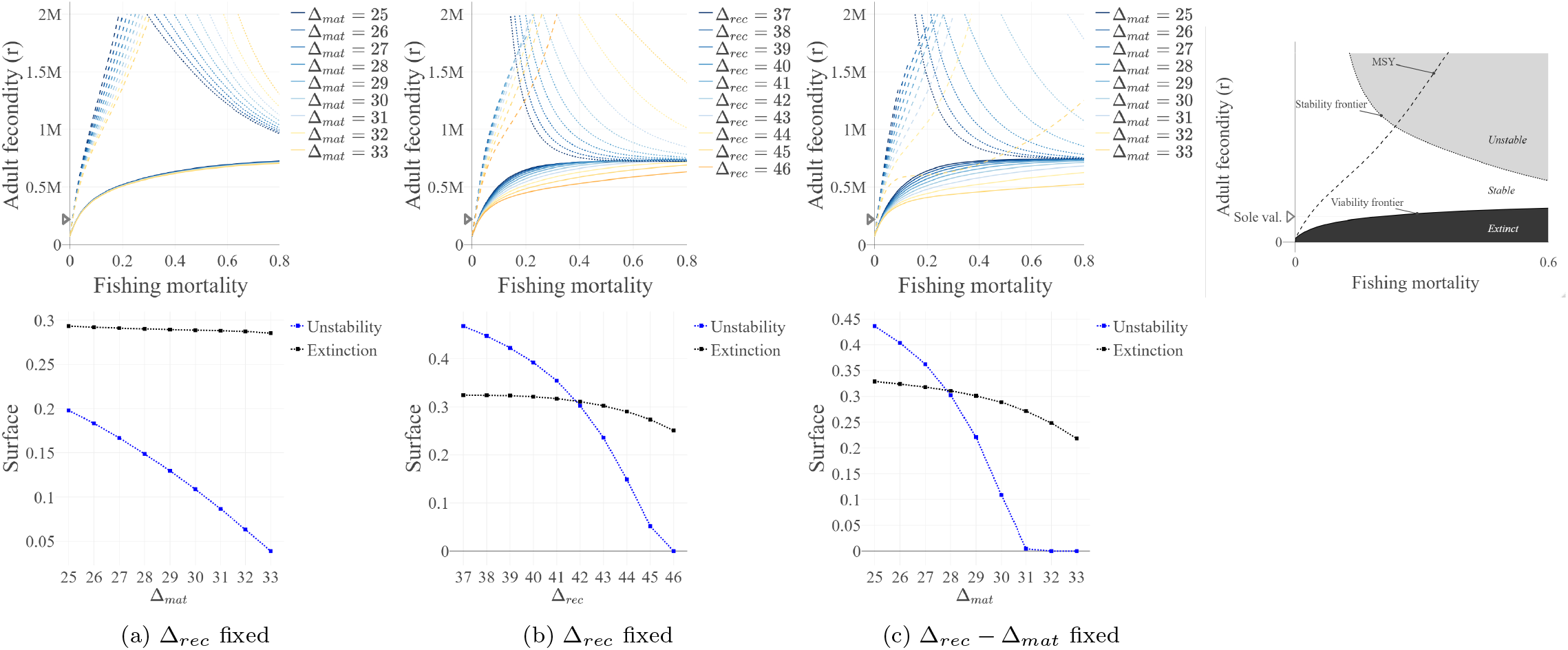
Stability of inter-annual equilibrium and viability domain, with associated surfaces (expressed as a proportion of the represented portion of the plan) in the *r* × *F* plane when Δ*_mat_* and/or Δ*_rec_* vary. All other parameters are set for the Bay of Biscay sole. For a given (Δ*_mat_*, Δ*_rec_*) pair corresponding to a colour shade, the solid line indicates the viability frontier, the dotted line indicates the stability frontier of the inter-annual equilibrium. The delimited zones are described schematically in the right panel. Finally, the dash line indicates the position of the *F_MSY_* for different values of *r*. The value of *r* for sole (*cf*. table 5) is indicated by the grey triangle. The three cases considered are: (a) Δ*_mat_* vary and Δ*_rec_* = 44; (b) Δ*_rec_* vary and Δ*_mat_* = 28; (c) both Δ*_mat_* and Δ*_rec_* vary and their difference is constant and equals 14 (Δ*_ma_t* = 25, …, 33; Δ*_rec_* = 39, …, 47).

When Δ*_rec_* only varies (figure 6b), as was seen before, an increase in Δ*_rec_* value increases the viability surface of the population. Indeed for a same value of parameter *r*, a higher *F* brings population to extinction when Δ*_rec_* is high. Moreover, as was stated above, a difference of one or very few months can have large consequences in terms of viability. This is especially true for species with large values of *r*.

Variations of Δ*_rec_* also have important consequences concerning the position of the stability frontier. As with Δ*_mat_* variations, for a given *r* the value of *F* necessary to destabilise inter-annual equilibrium is higher when Δ*_rec_* is high. This means that protection of mature individuals from fishing also has a stabilizing effect on the population.

As was also stated before, the value of *F_MSY_* is quite insensitive to variations of Δ*_mat_* and Δ*_rec_*, especially when *r* is low. By superimposing curves for *F_MSY_* with stability domain, we can see that MSY can in fact, at least theoretically, be associated to an unstable *i.e*. non-sustainable state. However considering realistic values for *r* (of the same order of magnitude as the sole of the Bay of Biscay, say 0.2M), it is clear that the population is much more likely to become extinct than to get his equilibrium destabilised, given the respective positions of the stability and viability frontiers (figure 6).

When Δ*_mat_* and Δ*_rec_* are varied jointly so that the mature unharvested phase is constant in duration (figure 6c), differences with the pattern observed for Δ*_rec_* (figure 6b) concerning stability and viability frontier are small. However, differences exist on *F_MSY_* curves. By contrast to the pattern observed for Δ*_mat_* variations only (figure 6a), they are interrupted when *r* is increased beyond a certain threshold. This interruption is due to a modification of the yield curve’s shape, the optimum being replaced by a plateau (see figure E.11 in appendices). In this configuration the MSY strictly speaking (*i.e*. the optimum) was always stable in the ranges of *r* and *F* considered.

In difference equations models, it is well known (Ricker, 1954) that population stability can be visualised by plotting the stock-recruitment relationship. In our model, the relation between the number of individuals participating to reproduction (*i.e* the “stock”) and the associated number of individuals entering the exploited portion of the population (*i.e*. the “recruitment”) emerges from density-dependent processes involving several classes of individuals and occurring at different time-steps (see Appendix D for the complete formulation of the relation between spawning stocks and associated recruitment). However, it is possible to generate observations of recruitment as a function of stock by running the model for a number of time steps with different initial conditions.

When the model is run with realistic parameters for sole (figure 7a), which corresponds to a region at the bottom-left of the parameter plan represented in figure 6 (indicated in this figure by the grey triangles), the observed relationship is monotonous and steadily increasing. It shows here no evidence of decreased recruitment for high values of stock. Observations are roughly arranged along a continuous curve which means that oscillations are quickly damped.

**Figure 7:**
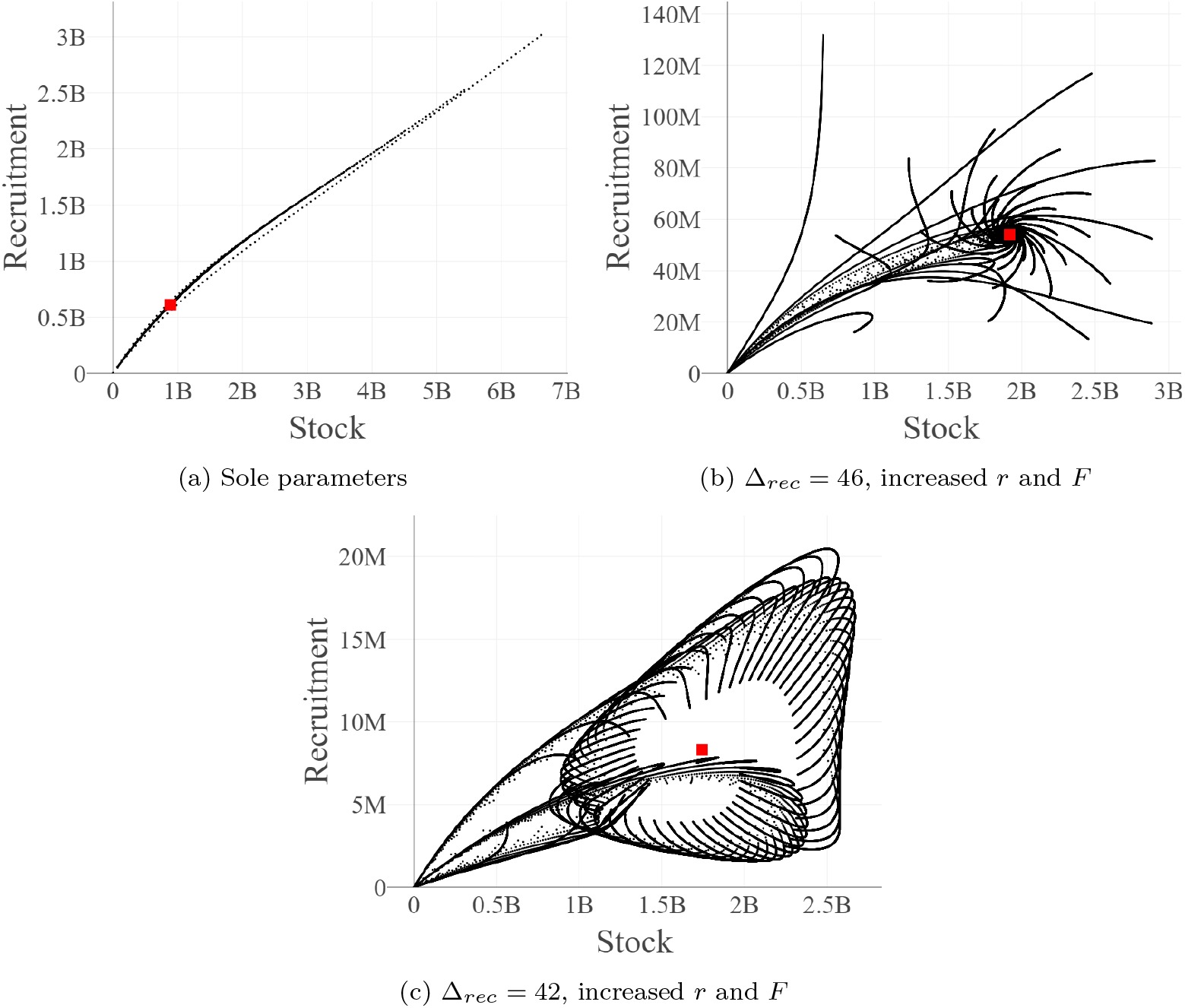
Stock-recruitment relationship extracted from simulated time-series of the model. Each black point represents an association of a given stock (number of mature individuals at *m_mat_*) with the corresponding recruitment (recruted inidividuals as defined in our model) after *a_rec_* years, at the same month *m_mat_*. Values are simulated by running the model for 1000 time steps with 1000 initial conditions evenly distributed between a stock of 10 and 10^10^ individuals. Equilibrium stock and recruitment are represented by the red square. Different sets of parameters are tested: (a) model parameterised for sole with Δ*_mat_* = 28 and Δ*_rec_* = 44, (b) same parameters but with *μ* and *ω* re-estimated for Δ*_rec_* = 46, *r* = 1.5 × 10^6^, *F* = 0.45; (c) same parameters but with *μ* and *ω* re-estimated for Δ*_rec_* = 42, *r* = 1.5 × 10^6^, *F* = 0.45

When we move to a higher region of *r* × *F* plan presented in figure 6, with *r* = 1.5 × 10^6^ and *F* = 0.45, we can observe more complex dynamical behaviours, as well as the destabilisation of equilibrium with a decreased Δ*_rec_*. In the case where Δ*_rec_* = 46 (figure 7b) for example, which corresponds to a stable region of the parameter space (see figure 6b) there is a concentration of points around the equilibrium value which indicates stability of the latter, even if large oscillations are observed before stabilisation. When Δ*_rec_* is decreased from 46 to 42, we move from a stable to an unstable region of the parameter space (see figure 6b). Then, we can observe no stock-recruitment pairs in the neighbourhood of the equilibrium (figure 7c) which is a clear sign of destabilisation of the equilibrium. Instead, we get very complex trajectories which indicate apparition of chaotic oscillations.

Although we observed that, especially for the two cases with increased *r* and *F*, a single value of stock could be associated to a set of possible recruitment values, we can recognise in the scatter-plots the apparition of the typical dome-shaped stock-recruitment relationship when moving from a very stable to a less stable and unstable region of the parameter space (figures 7a-7c).

### 3.5. Sensitivity of the resilience to timing of recruitment and maturing

Resilience also is affected by intra-annual variations of Δ*_mat_* and Δ*_rec_*. Once again, the influence of Δ*_rec_* is more pronounced than the influence of Δ*_mat_*. First of all it must be noticed that for most of the (Δ*_mat_*, Δ*_rec_*) pairs considered, we had *F_MiRT_* very close to 0 which means that resilience generally decreased with increasing *F*. Values of *MiRT* were also largely insensitive to variations of Δ*_mat_* and Δ*_rec_*. The very small variance of *MiRT* was mainly explained by variations of Δ*_mat_* (*SI* = 0.77) whereas the one of *F_MiRT_* was mainly explained by Δ*_rec_ (SI* = 0.85), as shown in figures 4e and 4f.

All resilience curves tended toward infinity when *F* increased. We found not much difference between resilience curves when Δ*_mat_* only was varied. On the contrary, an increase of Δ*_rec_* was associated to the vertical asymptote moving to the right (figure 8b).

**Figure 8:**
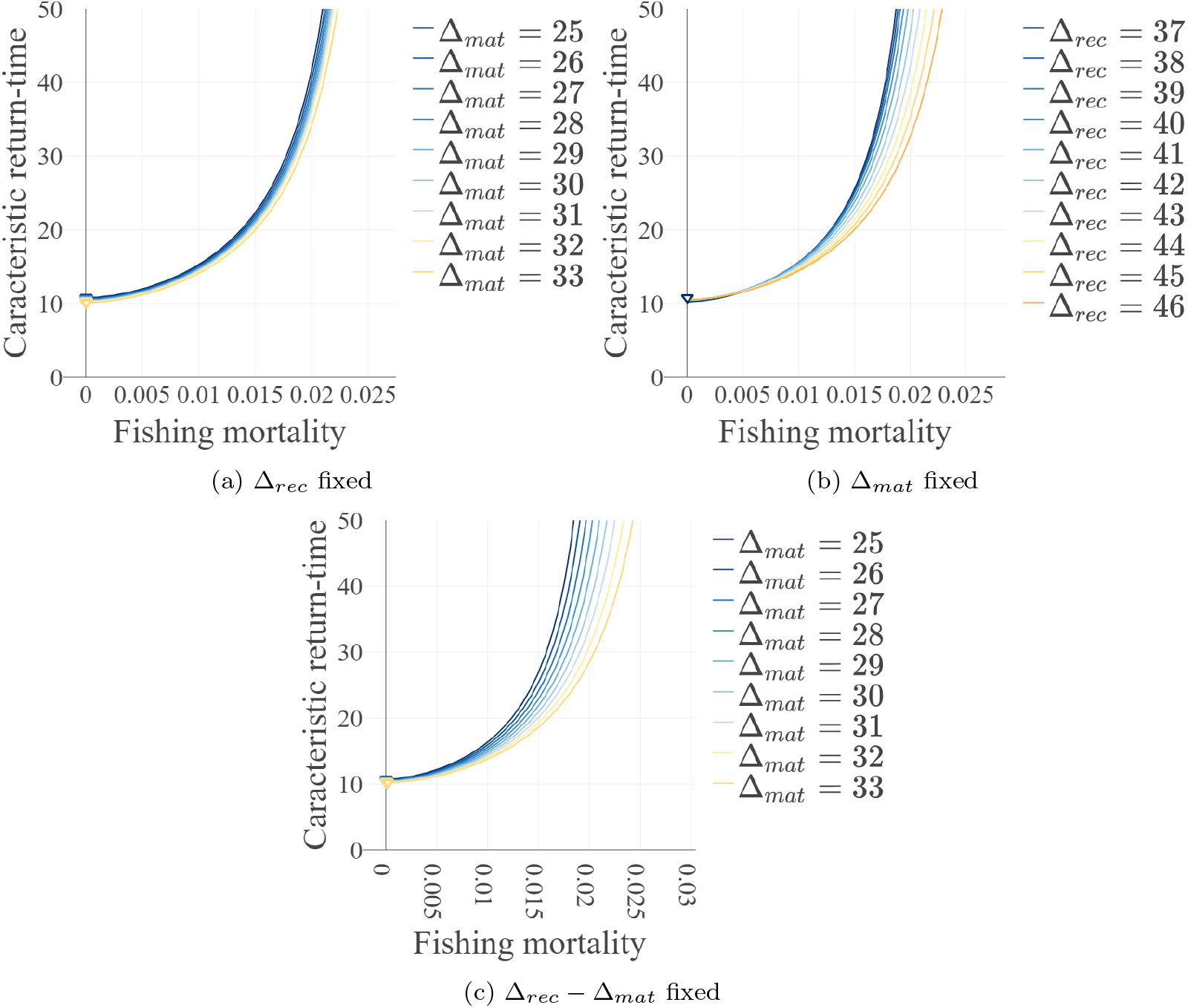
Caracteristic return-time to inter-annual equilibrium as a function of fishing mortality (by month) when Δ*_mat_* and Δ*_rec_* vary and the system is parameterized for the Bay of Biscay sole. Different colours indicate different values of Δ*_mat_* and/or Δ*_rec_* depending on the case: (a) Δ*_mat_* vary and Δ*_rec_* = 44; (b) Δ*_rec_* vary and Δ*_mat_* = 28; (c) both Δ*_mat_* and Δ*_rec_* vary and their difference is constant and equals 14 (Δ*_mat_* = 25,26, …, 33; Δ*_rec_* = 39,40, …, 47).

Combining the results to provide a biological interpretation for the Bay of Biscay sole (with r supposed to be near 0.2M, see Appendix D for details of parameters value), the loss of resilience observed when increasing *r* characterises the approach of the viability frontier rather than the stability frontier. The consequence of this observation is that it is possible to have population weakly resilient even far from the stability frontier if the viability frontier is near. Even when fishing at *F_MSY_* this situation is likely to be problematic when *F_MSY_* is near from the frontier, *e.g* for species with low *r* or high Δ*_mat_*.

## 4. Discussion and conclusion

Our aim in this study was to investigate the effect of processes occurring at intra-annual time scales on the amount and sustainability of long-term yields from a population, and on the widely used reference point known as MSY. We considered an idealised population submitted to three structuring processes namely reproduction, maturation and recruitment and described using a deterministic, structured dynamic population model in monthly discrete-time. The main originality of the modelling approach relies on the particular way we structured the population in order to (i) make the analysis tractable, (ii) link intra-annual and inter-annual time-scales and (iii) to avoid resorting to stock-recruitment relationship to represent the dynamics. Most modelling approaches use structuring by age (Marchal et al., 2009; Tahvonen, 2009; Doyen et al., 2012; Nielsen and Berg, 2014), by size (Bartolino, 2011; Lindstrøm et al., 2009) or by stage (Zipkin et al., 2008; Wikström et al., 2012; Liz and Pilarczyk, 2012). In our modelling framework instead, the classes of the population are defined by combining age (in year), maturity and accessibility to fishing characteristics. Thanks to this original structure we were able on the one hand, to simulate month by month the evolution of the population, and on the other hand, to resume the inter-annual dynamics to a first order difference equation system. This interesting result enabled us to compute analytically the inter-annual equilibrium and to assess numerically its stability and resilience as a function of the model parameters. This is in our sense one of the main innovation of our approach.

Application of the model was illustrated with parameters and data published for the Bay of Biscay sole, although we stress that our model is general and flexible enough to be applicable to any species whatever the ages of maturation and recruitment. In our model, we hypothesized that recruitment occurs after maturation but the reverse situation would constitute a straightforward generalisation.

First, we found that yields curves shape was influenced by variations of maturation lag Δ*_mat_* and recruitment lag Δ*_rec_* in different ways. The value of MSY was more sensitive to Δ*_mat_* whereas viability, stability and resilience were more sensitive to Δ_*rec*_. In classical stock-recruitment modelling (Hilborn and Walters, 1992), all processes occurring before the age of first capture are synthesised into a single stock-recruitment relationship (Bjorkstedt, 2000). The advantage of our model is to separate explicitly maturation and recruitment as processes of different nature. As a matter of fact, maturation is a strictly biological process on which no control is possible whereas recruitment is in part dependent of fishing behavior and gear (Laurec and Le Guen, 1981) so that it could be considered as a control variable. Simulations of the emergent stock-recruitment relationship in our model show that under certain conditions, this relationship can be complex and a single value of stock associated to a large number of potential recruitment values. These oscillations must be a consequence of the separation of the stock into different stages. Soudijn and de Roos (2017) found that adding juvenile stages in a population model enhanced population cycles and made dynamics more realistic in the sense that they approximated better a physiologically structured model. Exhaustive description of attractors associated to observed oscillations, although interesting, is beyond the scope of this study. It remains that this result outlines the interest of modelling explicitly life-history processes occurring in the youngest stages of the population. However, it is true that the most complex dynamical behaviours were obtained with values of fecundity probably unrealistically high. With realistic values of parameters, these quantities would have been quite well described by a classical stock-recruitment function.

In practice, for most exploited species, these information on maturation and recruitment are known, if at all, on a yearly basis only ICES (2018) and the specific uncertainty related to timing and duration of processes on intra-annual time-scales is generally ignored. The application of our model to the Bay of Biscay sole is in line with other studies (Kokko and Lindström, 1998; Tang and Chen, 2004; Xu et al., 2005; Cid et al., 2014) concerning the fact that intra-annual time-scales matter and that neglecting them can lead to important errors. Here, we argue that timing of biological processes and harvesting have different effects. With this example we show, on the one hand, that ignoring intra-annual timing of maturation can have a large impact on the computation of classical reference points such as MSY. On the other hand, we found that the viability frontiers of the population was sensitive to small variations in recruitment time. It is expected that these aspects could be even more critical if the seasonality of fishing was considered and F varied within the year as it is usually the case. Fortunately, the value of *F_MSY_* was insensitive to variations in Δ*_mat_* and Δ*_rec_* so that the advice for the population management at *F_MSY_* would not be much affected by uncertainty concerning processes occurring at intra-annual time-scales.

The second aspect of our study was to quantify local stability of inter-annual equilibrium as a measure of sustainability of yields drawn from the population. There is a growing debate on whether the fish populations are stable or not, and on the role of fishing on their destabilisation (Anderson et al., 2008; Shelton and Mangel, 2011; Sugihara et al., 2011; Rouyer et al., 2012). Our results support after other studies (Hsieh et al., 2006; Anderson et al., 2008; Cid et al., 2014; Liz, 2017) the fact that single population’s equilibrium can be destabilised by increasing fishing mortality, but only for species with very high fecundity. When parameterised for the Bay of Biscay sole, the value of parameter *r* required to effectively destabilise the inter-annual equilibrium is too high to be realistic and the population is more likely to become extinct than to have his inter-annual equilibrium destabilised.

Shelton and Mangel (2011) assessed the stability of a Ricker model parameterised for 45 exploited stocks and concluded that the presence of deterministic cyclic or chaotic behavior in real stocks was very unlikely. Our results are consistent with this prediction even if differences with their findings must be noticed. In particular, an important difference concerns the modification of the stability region when the time between birth and maturation increase. In our model, the stability region increases when Δ*_mat_* increase, in the sense that a higher *F* is necessary to destabilise the inter-annual equilibrium. In their study, on the contrary, the stability region decreases with each year added between birth and maturation. The explanation of this difference must rely on the different hypothesis concerning density dependence. Indeed, in our model immature individuals’ mortality is made dependent of adult abundance at each month and not only on the abundance at birth-time as it is the case in Ricker model. Unfortunately, our model is too complex to get an analytical demonstration of this difference.

Our results demonstrate that the MSY can theoretically imply an unstable inter-annual equilibrium. This is in line with results obtained by Kokko and Lindström (1998). This eventuality was also known in predator-prey models (Beddington and Cooke, 1982) and in single species with Allee effect (Ghosh et al., 2014). Then we agree with Beddington and Cooke (1982) that sustainability of MSY reference points should not be taken for granted but we temper this view by saying that the risk to get an unstable MSY in the mono-specific case is very low. However, even in case of a stable equilibrium, resilience measured by the return time to this equilibrium should still be considered. Indeed, as stated Beddington et al. (1976), in some cases, *“perturbations may take so long to die away that effectively the populations may never return to equilibrium within a biologically meaningful times-pan”*. Such a situation would result in the impossibility of managing properly an exploited system.

Assessment of resilience in exploited populations is a topic of growing interest among empirical (Britten et al., 2014; Mumby et al., 2016) and theoretical ecologists. Tromeur and Loeuille (2017) and Kar et al. (2019) investigated relationship between the objectives of resilience and yields in food chains and found a RMY distinct from the MSY, leaving room for a trade-off between these objectives. Lundström et al. (2019) also explored trade-offs between yields and a number of conservation objectives including resilience on two structured single population models. One of their key result is that resilience is highly correlated with biomass loss, suggesting to use this metrics as a proxy for resilience in practice. This is in line with our observation that return-time to inter-annual equilibrium increase dramatically near the viability frontier. Here, we predict that in most real exploited populations, resilience will be harmed by approaching this frontier due to a lowered equilibrium biomass, rather than by getting in an unstable region of parameter space. In the theoretical studies above cited, values of parameters were set arbitrarily in an exploratory purpose and their authors found an optimum of resilience corresponding to a non-zero value of fishing mortality. Here, with a set of realistic parameters, we located this optimum at very low, although non zero, values of *F*. This would practically exclude some “win-win” situation in which it would be possible to increase both yield and resilience.

Previous works proposed to manage the trade-off between yields and resilience by acting on the distribution of effort on trophic level (Tromeur and Loeuille, 2017; Kar et al., 2019) or on the stage development (juvenile *vs*. adults) inside the population (Lundström et al., 2019). Our model is not designed to answer these questions. Instead, we evaluated the effect of another control parameter which is the timing of recruitment. In our application for the Bay of Biscay sole, we found that variations in recruitment time had a non-negligible effect on resilience curves although it did not affect much the value and location of the optimum.

Our aim in this study was to propose a generalised and relatively simple model to shade light on the effect of intra-annual time-scales in maturation and recruitment processes which are of key importance in fisheries management. Although we parameterised the model with published literature and data concerning a real stock, this was on a qualitative and illustrative purpose rather than to make quantitative predictions (*i.e*. to get possible rather than exact values). We stress that our model is far too idealised to make precise predictions and that available data are not designed for our model.

Our choice here was to formulate as simply as possible a model of density dependence by cannibalism of adults on juveniles. Cannibalism is known to be frequent in fish populations (Smith and Reay, 1991) including in highly exploited stocks such as cod (Bogstad et al., 1994; Uzars and Plikshs, 2000). Moreover cannibalism was at the core of development of Ricker historical stock and recruitment theory (Ricker, 1954) and is a useful assumption in the sense that it is the most straightforward process of over-compensatory mortality. Other processes such as competition for food and space (Biro et al., 2003) are known to potentially give rise to the same type of mortality. Rindorf et al. (2022) gave support to Ricker’s (1954) assumptions by showing that most of exploited stocks undergo density-dependence before recruitment and that overcompensation was more likely to occur in demersal stocks such as the Bay of Biscay of sole. From a dynamical point of view, over-compensatory reproduction curves with inter-stage density-dependence are known to cause more complex behaviours. We were interested in stability and resilience properties of the population equilibrium and for that reason adopting a number of Ricker’s assumptions was a way of not restricting the dynamical potential of our model.

This is to our knowledge the first resilience analysis on a single exploited population subject to cannibalistic density-dependence, as well as the first evaluating the effect of intra-annual timing of maturation and recruitment on resilience. Although we restricted ourselves to a particular case study and made a number of assumptions that limit the generality of the presented results, we stress that in future studies, the model could be modified in order to address new questions while adopting a similar approach for structuring the population in order to be able to study its dynamics easily as a first order system. For example, instead of dealing with the case in which maturation is separated from recruitment (what we did here), one could write another slightly different model in which start of reproduction is separated from end of density-dependent mortality. Alternatively, the reverse case where recruitment occurs before maturation would be worth exploring given that it would be verified in some species (ICES, 2018). All these extensions are permitted by the conceptual framework presented in this study.

Moreover, in an operational purpose, one could be interested in expanding this theoretical model to increase realism, at the cost of simplicity and analytical tractability. A natural extension to it would be to add a representation of fecundity depending on length or weight of individuals. This function would probably be species-dependent. Therefore, to maintain a broad scope of conclusions, it would be necessary to explore a large panel of functions. Also, the processes such as reproduction, recruitment and maturation were considered instantaneous while they are likely spread over several months in real populations. Finally other formulation for density-dependence including form of functional response in the cannibalistic case could be tested.

More generally, the absence of consideration of process error (*i.e*. error arising from under-specified models) limits the scope of our results. Here, we presented some non-linear dynamics obtained assuming perfect knowledge of the underlying mechanisms, but it remains true that non-linearities can be enhanced in models containing process error when this error propagates in a specific way (Anderson et al., 2008).

Our results suggest that ignoring intra-annual dynamics would result in little error on *F_MSY_* advice. However, we saw that small variations of the recruitment time would have non negligible consequences on the fishing mortality a population can support before extinction as well as on her resilience. We recommend that the latter be taken into account in harvest management, so that the sustainability of yields be guaranteed. A first step could be to modify the *F_MSY_*-range framework (Hilborn, 2010; Rindorf et al., 2017) to include resilience objectives to be attained aside from high enough yields.

## Appendix A. Alternative case, when *m_mat_* > *m_rec_*

The same way we deduced system (17) from equations (3–10) for the case when *m_mat_* ≤ *m_rec_*, we can deduce from the same equations the system E.1, expressed in table E.8. As before, this system can be expressed as a first order difference equation (see Appendix B for proof):

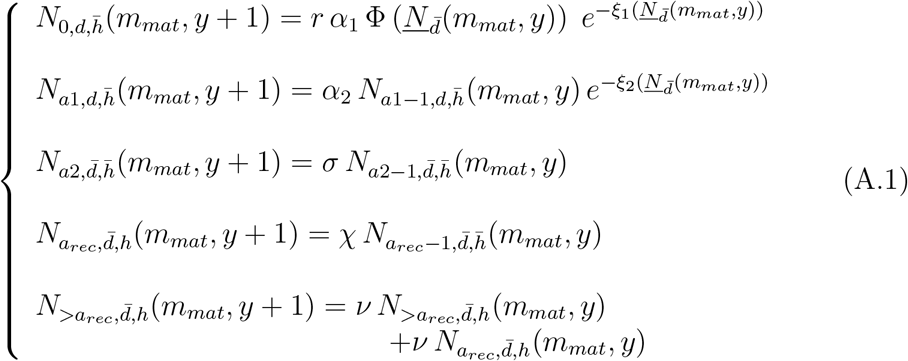

with *a*1 = 1, …, *a_mat_* and *a*2 = *a_mat_* + 1, …, *a_rec_* – 1, 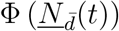 the number of spawners, 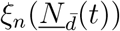 a density-dependence function. Here, we have :

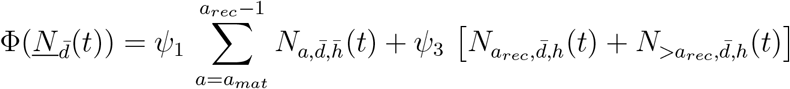

and

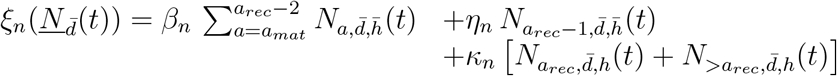

where

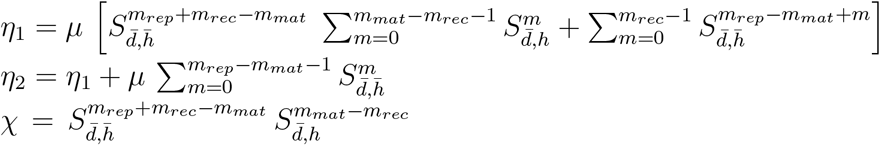

is a constant and all other parameters are the same as in model (11) (see table 4 for full detail).

The inter-annual equilibrium is solution of :

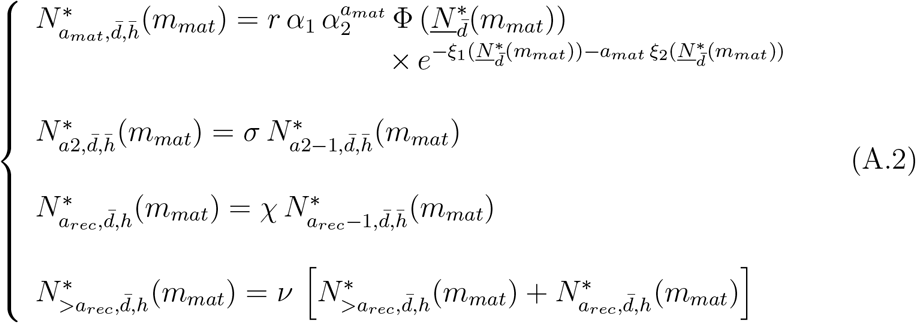

with

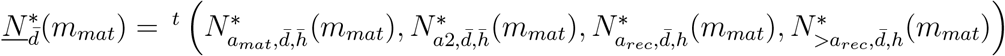

and *a*2 = *a_mat_* + 1 …, *a_rec_* – 1.

## Appendix B. Proof of result (11) and (A.1)

Expressions of systems (17) and (E.1) were deduced directly from the month model represented by equations (3–10). To rewrite these two systems as first-order difference equations systems we need to express:

1. 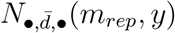
2. 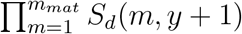
3. 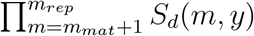

as functions of 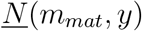.

### Expression of 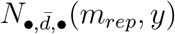

For all *y*, we have: 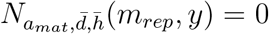 and 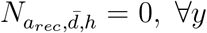. Hence, if *m_mat_* ≤ *m_rec_*, we get after equations (3–10):

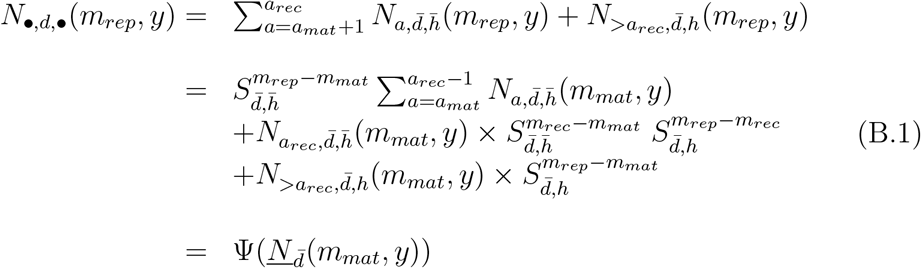

Else, we get:

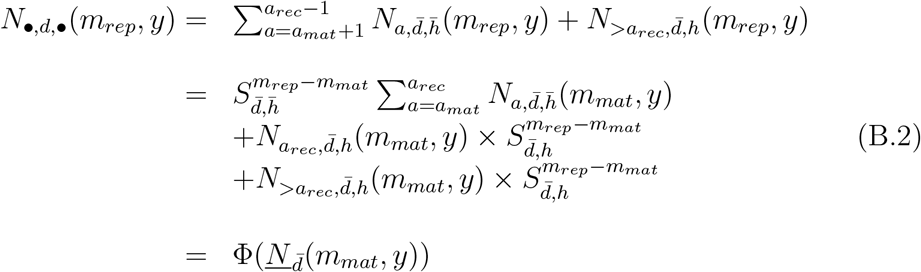

### Expression of 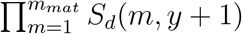

We have, for all *y*:

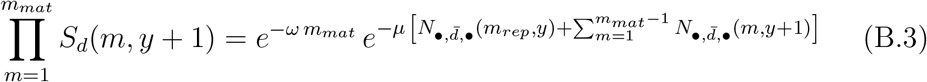

with

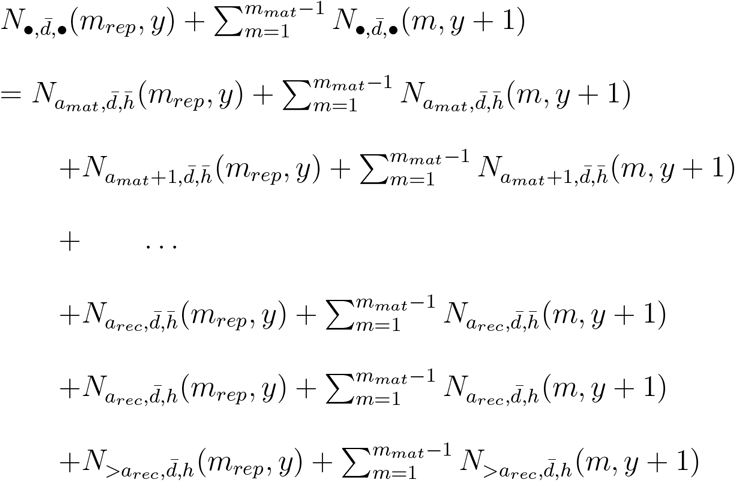

Moreover, after (3–10), we have for all *y* :

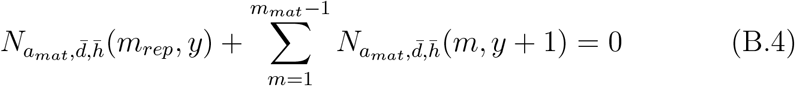

and for all *a* ∈ {*a*_mat_ + 1, …, *a_rec_* – 1}:

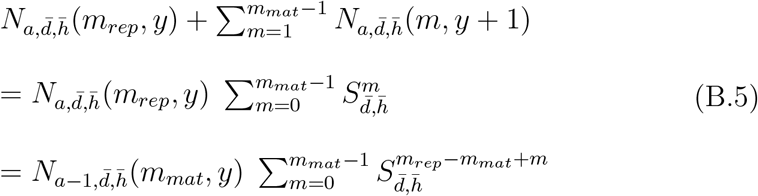

If *m_mat_* ≤ *m_rec_* we also have:

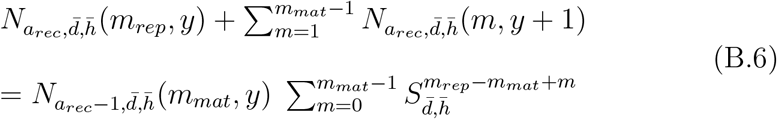

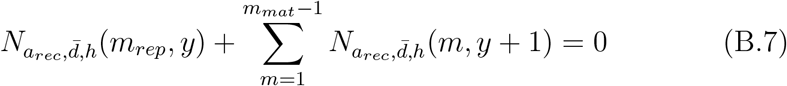

and

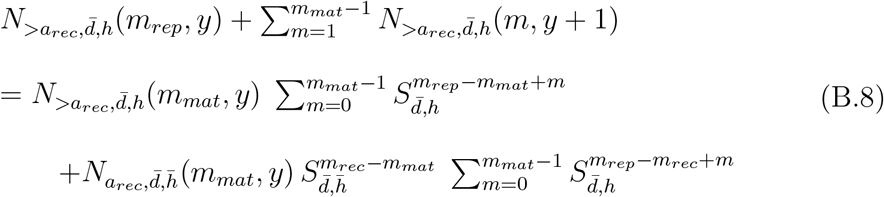

Else we have:

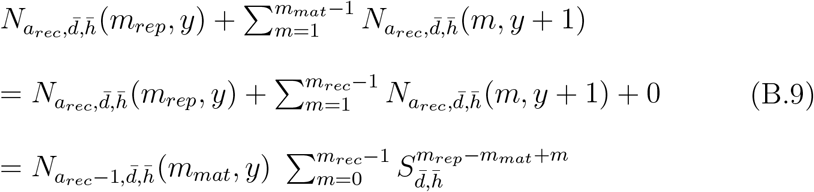

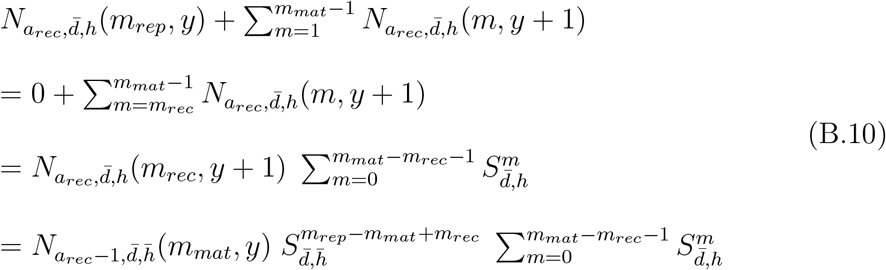

and

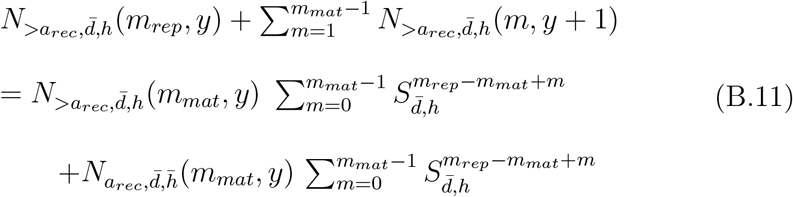

By summing equations (B.4–B.5) and (B.6–B.8) if *m_mat_* ≤ *m_rec_*, or (B.4–B.5) and (B.9–B.11) otherwise, and injecting it into (B.3), we can rewrite the product of interest as:

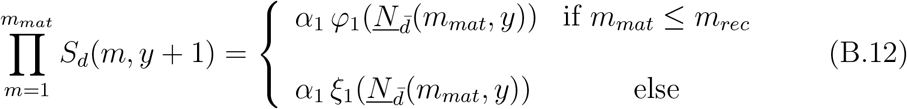

see table 4 for exact formulation of *α*_1_, *φ*_1_ and *ξ*_1_.

### Expression of 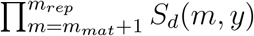

Likewise, we can express 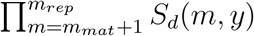 for all *y* as:

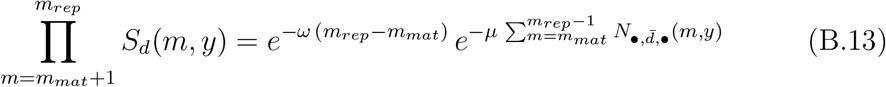

with

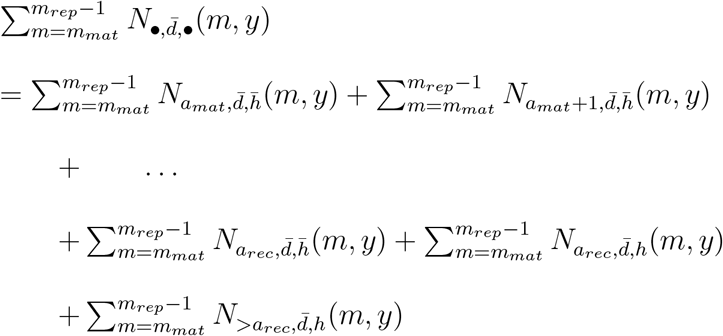

For all *a* ∈ {*a_mat_*, …, *a_rec_* – 1}, for all *y*, we have:

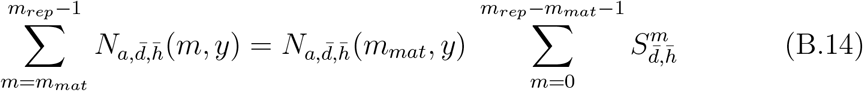

and

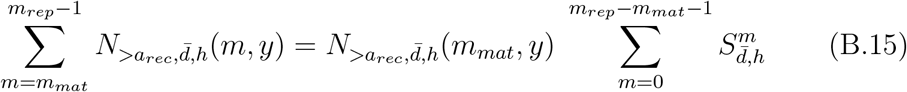

If *m_mat_* ≤ *m_rec_* we also have, for all *y*:

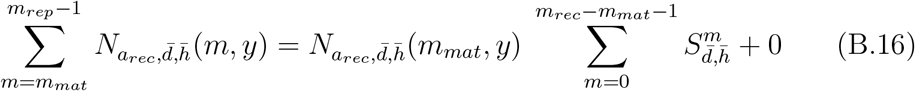

and

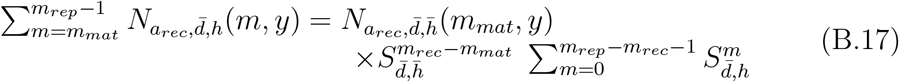

Else we have:

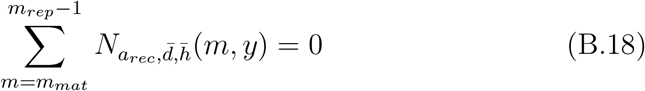

and

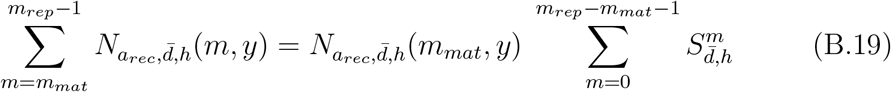

Hence, with notations exposed in table 4, we can write the double product:

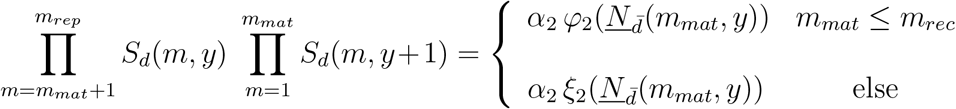

Finally we get systems (11) and (A.1).

## Appendix C. Jacobian of systems (11) and (A.1)

Let *J*\* (*m_mat_*) be the Jacobian matrix of system (11) at equilibrium. *J*\* (*m_mat_*) is written:

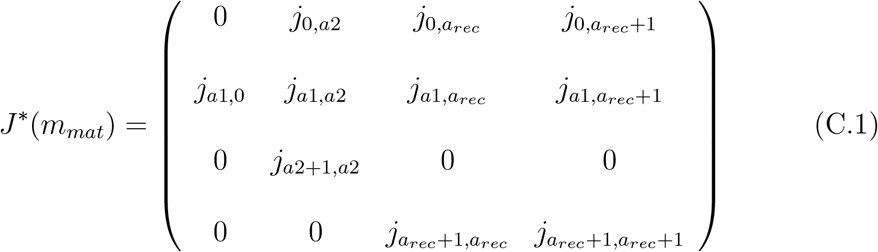

with *a*1 = 1, …, *a_mat_*; *a*2 = *a_mat_*, …, *a_rec_* – 1 and:

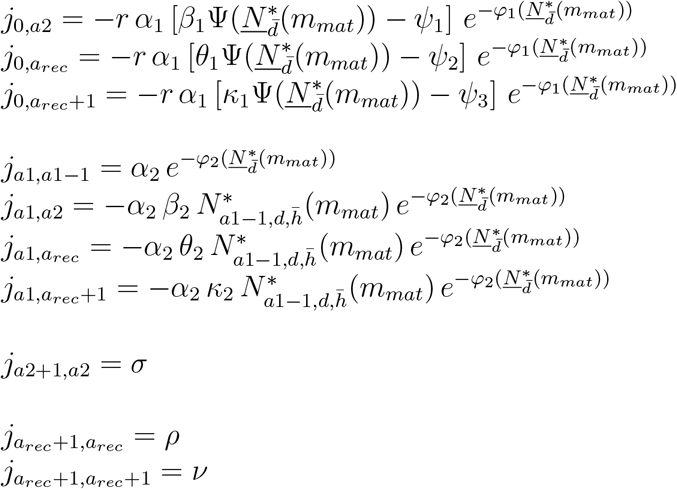

For system (A.1), the Jacobian matrix at equilibrium is of the same general shape but elements are instead:

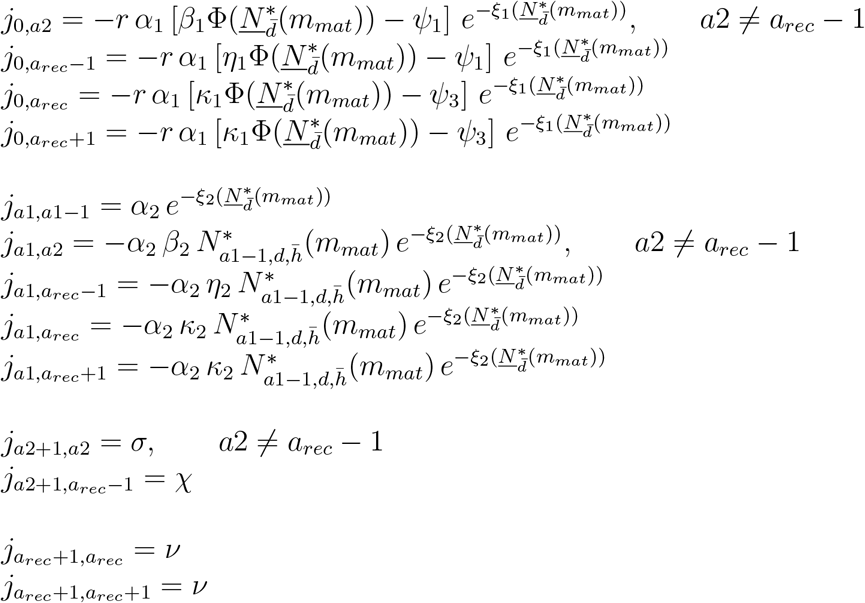

See table 4 for formulations of all the constants.

## Appendix D. Estimation of parameters for the Bay of Biscay sole

Assume that all the model parameters are known except *μ* and *ω*. Here, we aim at fitting a custom stock-recruitment relationship compatible with our model formulation to estimate those parameters. In the case where Δ*_mat_* and Δ*_rec_* verify *m_mat_* ≤ *m_rec_*, the population is governed by system (11). It comes immediately from the formulation of the system that:

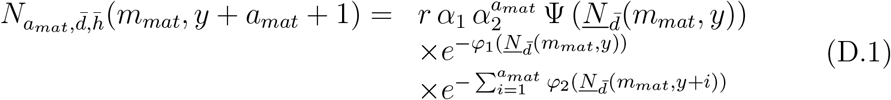

which is the number of newly mature individuals as a function of the set of mature individuals of the *a_mat_* years before.

To get the number of newly recruited individuals, we just need to take into account the years between maturation and recruitment. We get:

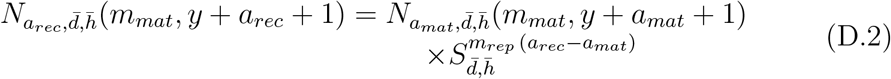

Let us pose: 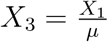 and 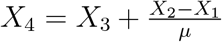, where *X_n_* = *β_n_, θ_n_, κ_n_* (see table 4 for computations of these constants), and let us define:

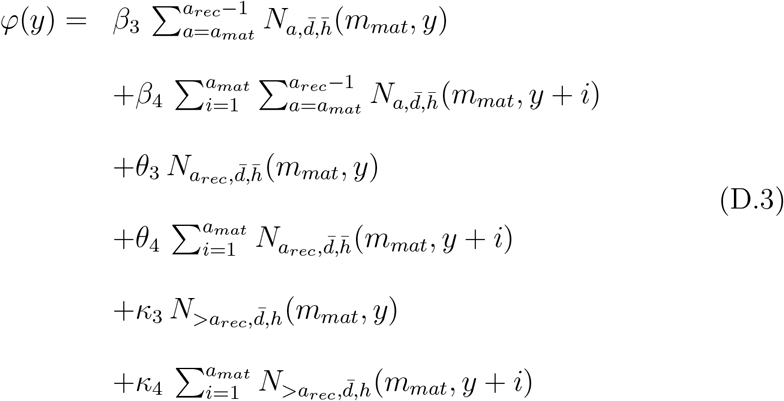

then we can express:

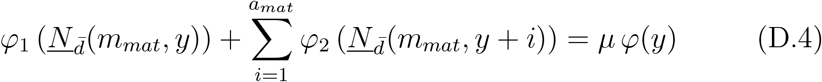

Finally, with 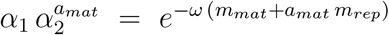 we can write the custom stock-recruitment relationship as:

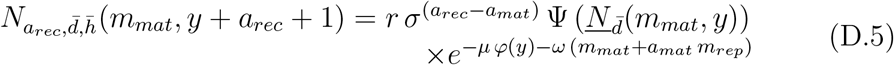

Conversely, in the case where Δ*_mat_* and Δ*_rec_* verify *m_mat_* > *m_rec_*, the relationship to be fitted is:

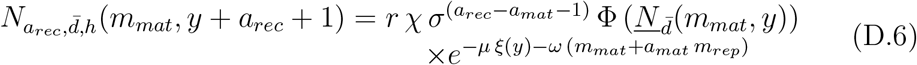

where

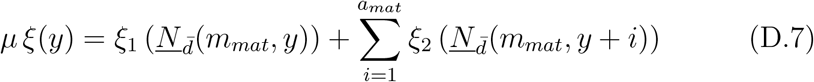

Hence, one can estimate parameters *μ* and *ω* by non-linear regression for any value of Δ*_mat_* or Δ*_rec_*, as long as sufficient data is available, assuming that assessment is effectively made at *m* = *m_mat_*. The regression was performed using the nls function of R software, and repeated each time Δ*_mat_* or Δ*_rec_* was modified.

**Figure E.9:**
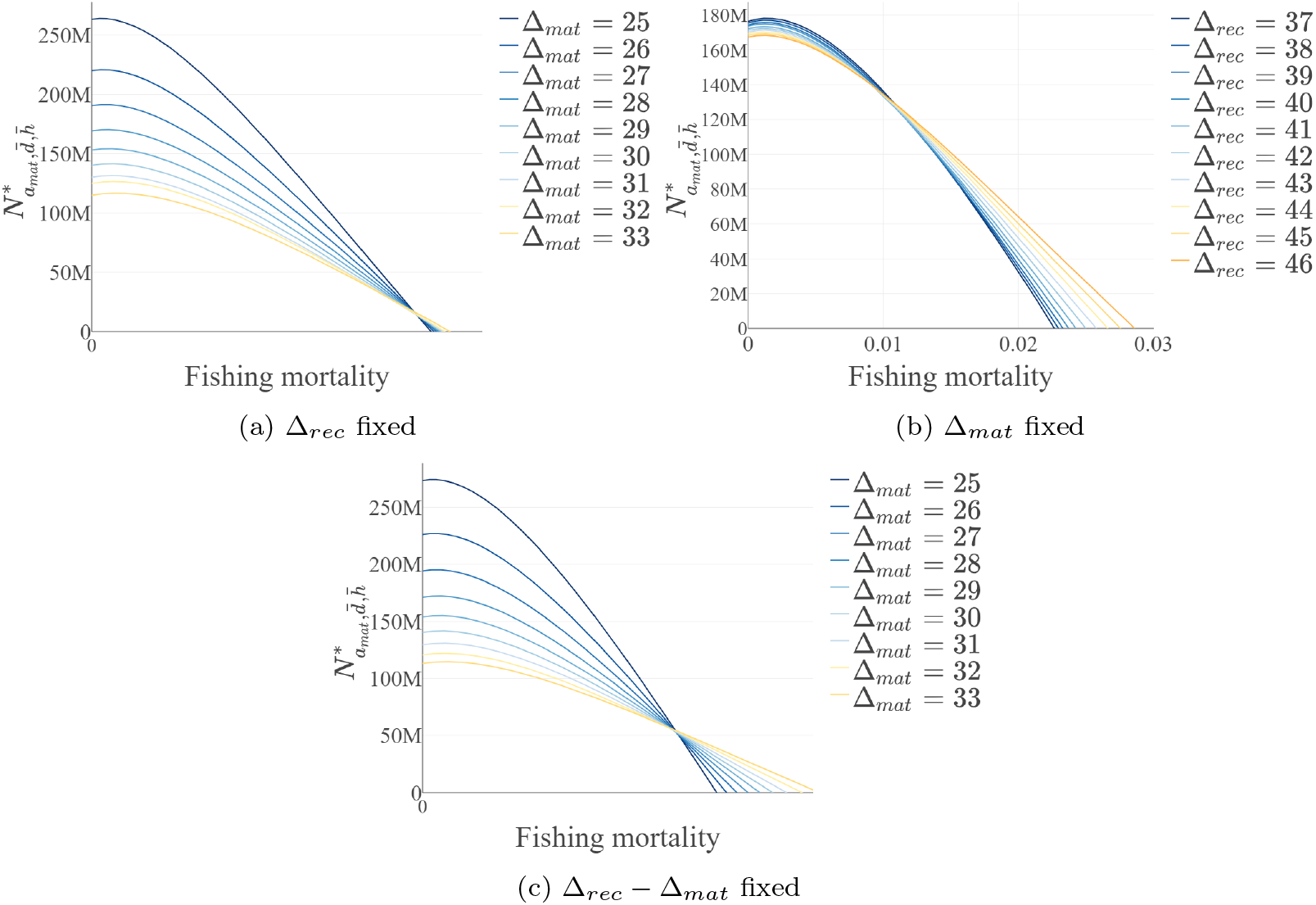
Number of mature individuals of round age *a* = *a_mat_* at equilibrium (*m* = *m_mat_*) as a function of fishing mortality (by month) when Δ*_mat_* and Δ*_rec_* vary and the system is parameterized for the Bay of Biscay sole. Different colours indicate different values of Δ*_mat_* and/or Δ*_rec_* depending on the case: (a) Δ*_mat_* vary and Δ*_rec_* = 44; (b) Δ*_rec_* vary and Δ*_mat_* = 28; (c) both Δ*_mat_* and Δ*_rec_* vary and their difference is constant and equals 14.

Estimations of parameters *μ* and *ω* for each combinations of Δ*_mat_* and Δ*_rec_* considered are given in tables 6 and 7.

NB: often, data for age *a_mat_ ≤ *a* < *a_rec_** are not available. In this case, one can compute them by applying the right constant to the first known age class. That is what we did for the Bay of Biscay sole (with *a_rec_ – a_mat_* = 1), considering that for all 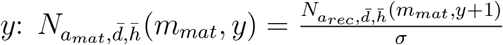.

## Appendix E. Supplementary outputs

**Figure E.10:**
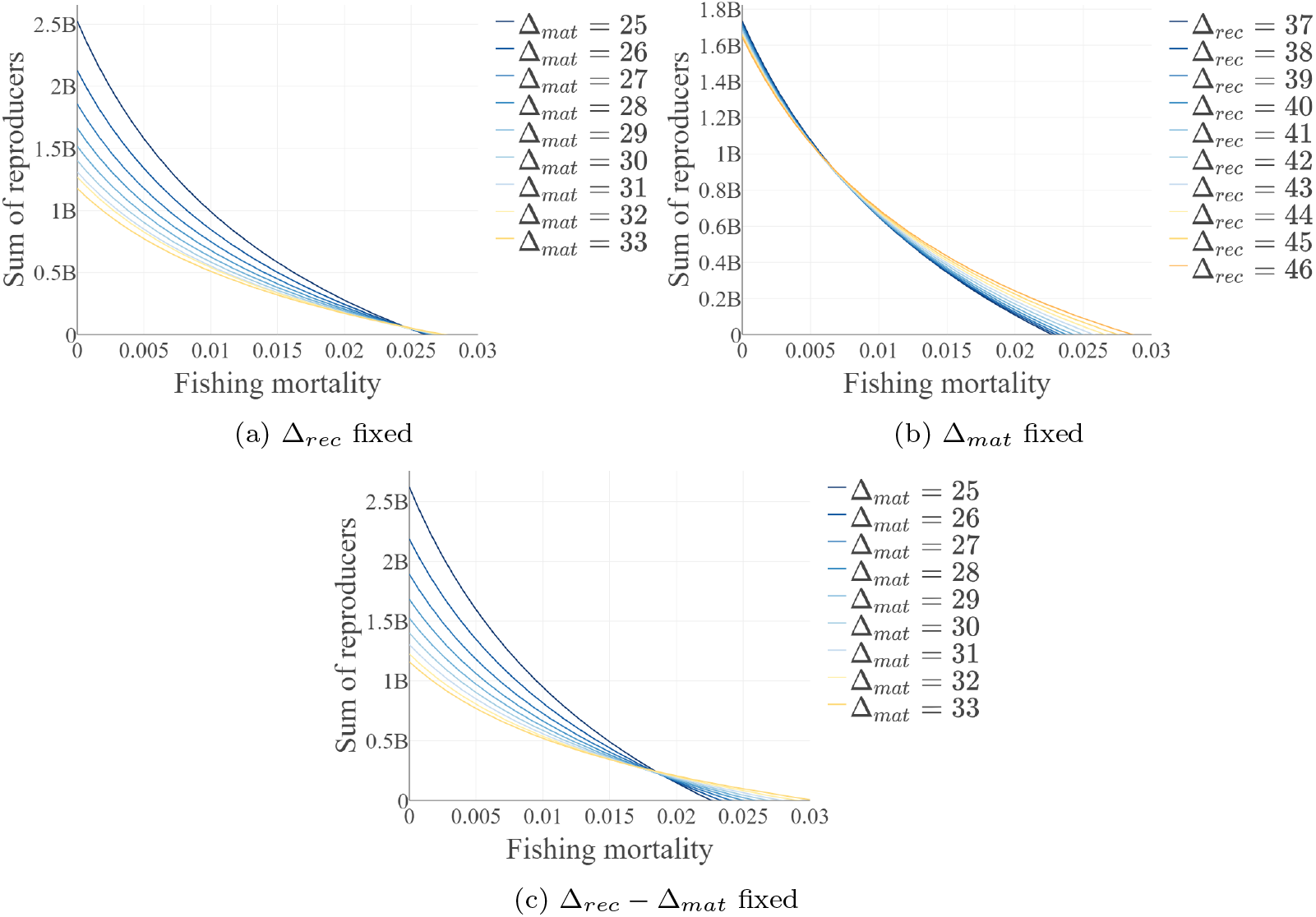
Sum of reproducers at equilibrium at *m* = *m_rep_* as a function of fishing mortality (by month) when Δ*_mat_* and Δ*_rec_* vary and the system is parameterized for the Bay of Biscay sole. Different colours indicate different values of Δ*_mat_* and/or Δ*_rec_* depending on the case: (a) Δ*_mat_* vary and Δ*_rec_* = 44; (b) Δ*_rec_* vary and Δ*_mat_* = 28; (c) both Δ*_mat_* and Δ*_rec_* vary and their difference is constant and equals 14.

**Figure E.11:**
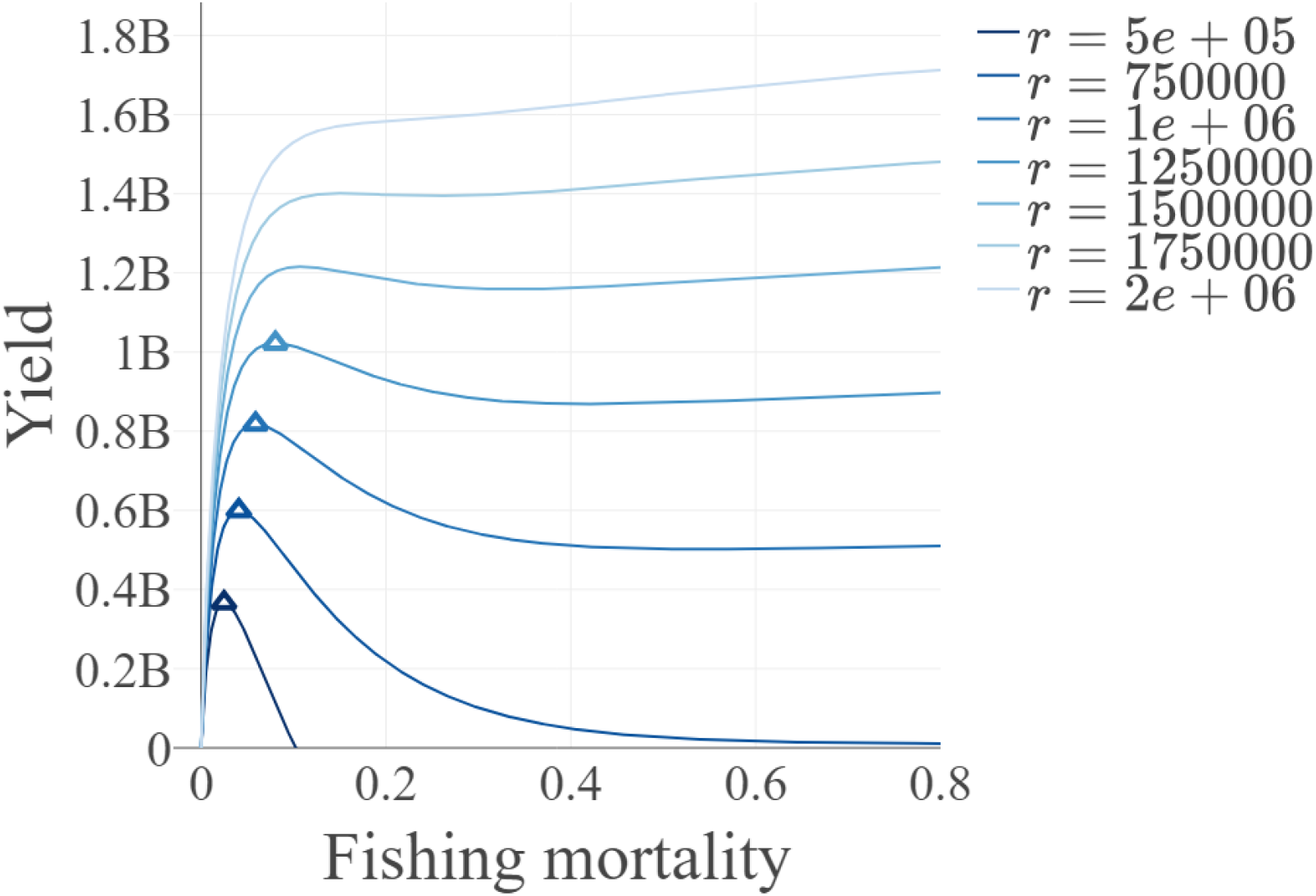
Total annual yields at inter-annual equilibrium as a function of fishing mortality (by month) and position of MSY when *r* varies, Δ*_mat_* = 25 and Δ*_rec_* = 39 (all other parameters being the same as in table 5). Coloured triangles correspond to MSY (numerically solved) and different colours indicate different values of *r*.

**Table E.8.**
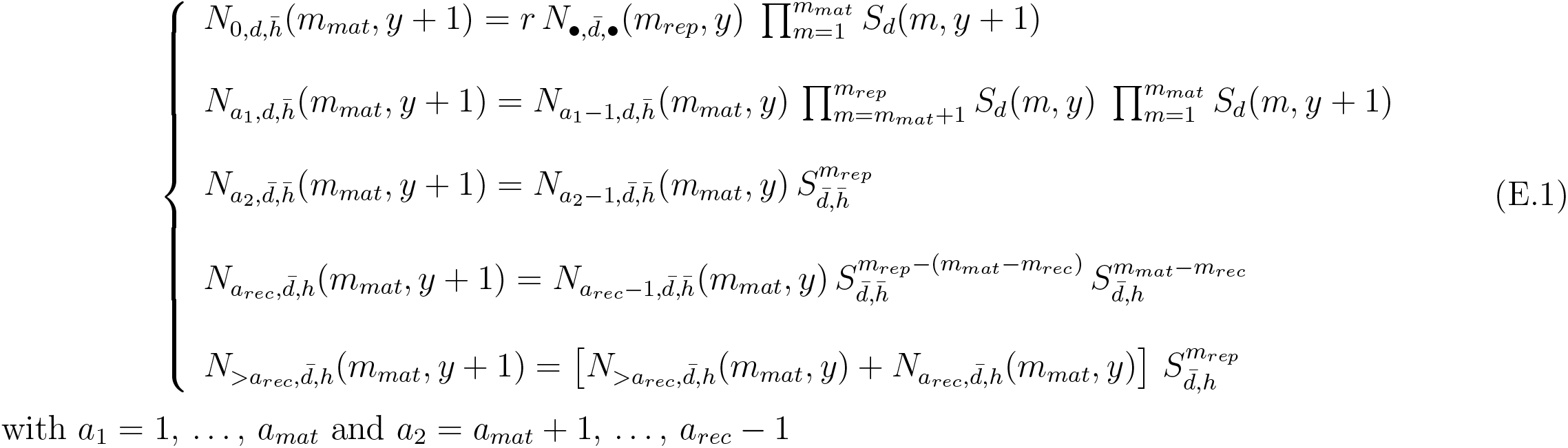
Expressions of the inter-annual dynamics of *<u>N</u>*(*m_mat, y_*) when *m_mat_* > *m_rec_*:

